# Intrinsic promoter responsiveness dictates sensitivity to transcriptional activation by enhancers

**DOI:** 10.64898/2026.06.25.734173

**Authors:** Yingxuan Tan, Judhajeet Ray, Maya U. Sheth, Benjamin R. Doughty, William J. Greenleaf, Jesse M. Engreitz

**Author notes:** Correspondence (W.J.G.), (J.M.E.).

## Abstract

Enhancers activate specific target promoters, but whether intrinsic enhancer-promoter compatibility contributes to this specificity is debated. Recent studies using different reporter assays have reached contradictory conclusions. We compare six reporter assay designs, identify confounders that bias compatibility measurements, and apply improved assays to test 25,000 enhancer-promoter pairs. Promoters differ in their responsiveness to enhancers (>100-fold versus 1.1-fold activation) while enhancers activate all promoters in a similar rank order. Promoter output scales with enhancer activity following a power law, with the exponent varying across promoters. Incorporating this exponent into the Activity-by-Contact model improves prediction of endogenous enhancer effects, explaining why certain active genes are insensitive to distal perturbations and “skipped” by enhancers. Responsiveness is modulated by transcription factor motifs in the core promoter. This work establishes responsiveness as an intrinsic promoter property that enables specific promoters to be highly activated in a landscape of broadly compatible enhancers, while others remain unaffected.

## Introduction

A fundamental question in human gene regulation is how enhancers activate specific target promoters in the genome. Genomic perturbation experiments have revealed complex patterns of enhancer-gene regulatory interactions, such as enhancers “skipping” over a nearby gene to regulate a more distant one ^1–5^. A long-standing goal has been to determine which factors influence enhancer-promoter specificity and how we can model these factors to predict the effects of enhancers on target genes (reviewed in ^6,7^). Existing predictive models of enhancer-gene regulation, such as the Activity-by-Contact (ABC) model and deep learning-based sequence-to-omics models, cannot yet explain many such complexities^1,8–12^. Resolving these questions is crucial for building predictive models of gene regulation in the genome ^8–11,13^, understanding molecular mechanisms of enhancer-promoter specificity ^14–16^, interpreting non-coding human genetic variation ^17,18^, and designing regulatory DNA for gene therapies ^17,19,20^.

One explanation for specific patterns of enhancer-gene regulation is intrinsic, sequence-based preferences between enhancers and promoters ^21–25^. Such preferences could be related to biochemical interactions or dependencies between transcription factors or cofactors at the enhancer and promoter ^6,26–29^. To interrogate these properties, a canonical experiment has been to clone combinations of enhancer and promoter sequences adjacent to one another in a reporter assay and measure their transcriptional activity (**Fig. 1A)**. Such reporter assays were used to discover the first transcriptional enhancers ^30,31^, and early studies suggested that enhancers were broadly promiscuous—capable of activating diverse promoters when placed next to each other on a plasmid ^32^. Over the past decade, massively parallel reporter assays (MPRAs) have expanded the scale of such studies and tested thousands of enhancer-promoter combinations at once in a variety of episomal and genomic contexts^27,33–38^. However, studies conducted in different systems using different assay designs have reached sharply different conclusions. Three distinct patterns of enhancer-promoter compatibility have been proposed (**Figs. 1B-D**):

**1. Broad/equal compatibility:** Each enhancer activates all promoters by the same fold-change – *i.e.,* an enhancer that activates Promoter A fivefold activates Promoter B fivefold as well (**Fig. 1B**). This pattern can be evaluated by modeling the expression level of a given enhancer-promoter pair as the basal strength of the promoter multiplied by the intrinsic activity of the enhancer (*Expression* ∼ *P_basal_* × *E*)^33^. This pattern has been supported by measurements of tens of thousands of enhancer-promoter pairs in various human cell lines^1,34^ and is qualitatively consistent with long-standing observations that many enhancers are compatible with various promoters in heterologous reporter systems ^32^.
**2. Promoter responsiveness:** Enhancers can activate all promoters, as in broad compatibility, but promoters differ in their intrinsic capacity to respond to activation by any enhancer. Enhancers produce the same rank order of effects across promoters but activate them by different magnitudes – *i.e.*, a strong enhancer might activate Promoter A tenfold but Promoter B by only twofold (**Fig. 1C**). Such differences in promoter responsiveness have been observed to varying degrees with small numbers of representative promoters comparing housekeeping and cell-type–specific genes in plasmid reporters ^36,39^ and genome-integrated reporters ^37,38^. One study observed a particular form of responsiveness in which expression of an enhancer-promoter pair depended on enhancer activity raised to a promoter-specific exponent (*Expression* ∼ *P_basal_* × *E^Presponsiveness^*)^36^.
**3. Combinatorial specificity:** Certain combinations of enhancers and promoters have intrinsic preferences for one another that cannot be explained by the above models – *i.e.* one enhancer activates Promoter A by tenfold and Promoter B by twofold while another enhancer activates Promoter A by only twofold but Promoter B by tenfold (**Fig. 1D**). Such preferences could take many forms and would need to be modeled using an interaction term specific to each enhancer-promoter pair. A striking example of such specificity comes from a study in Drosophila S2 cells, in which core promoters of developmental versus housekeeping genes were both highly inducible by enhancers but showed >10-fold differences in their activation by different classes of enhancers ^27^. In mammals, a study in mouse embryonic stem cells also observed strong specificity, finding that 60% of tested candidate CREs exhibited significant promoter selectivity for enhancers ^35,36^.

**Figure 1.**
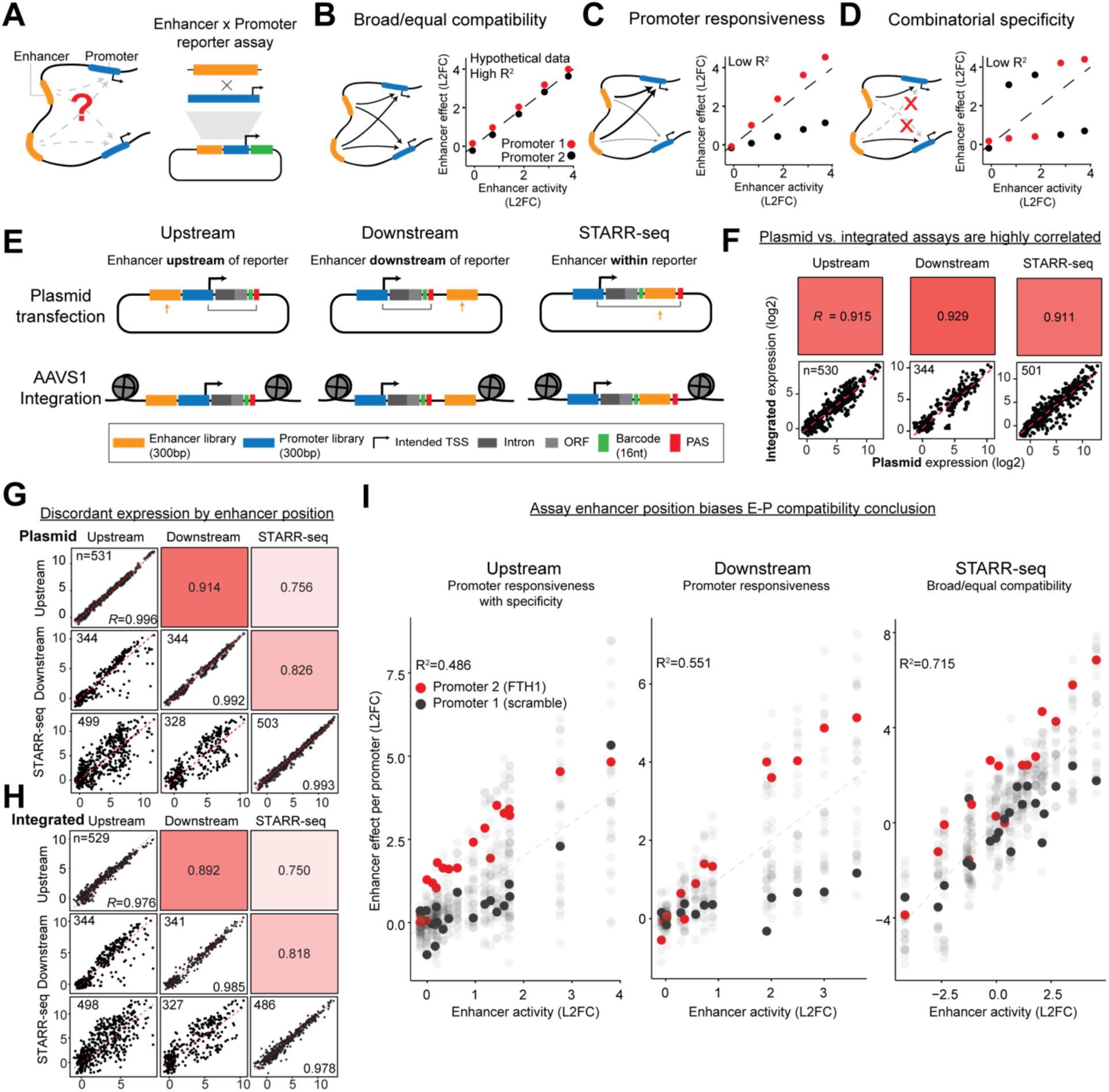
Enhancer location, not plasmid transfection, drives discordant E-P expression across different MPRAs. **A**, Combinatorial reporter assays for quantitatively assessing E-P compatibility. **B**-**D**, Proposed patterns of E-P compatibility (broad compatibility (**B**), promoter responsiveness (**C**), combinatorial specificity (**D**)) illustrated with hypothetical data from two promoters and a series of enhancers of varying strength. Each point represents an E-P pair while the axes compare enhancer effects on individual promoters (log2FC, y-axis) with enhancer activity (average log2FC effects, x-axis). Each enhancer has a single value for enhancer activity (*E_avg_*, x-axis) but multiple promoter-specific effects (one log2FC per paired promoter, y-axis). Vertical variance about the y = x line indicates unequal enhancer effects on paired promoters. **E,** Schematic of three common enhancer MPRA formats test in two different chromatin contexts. **F**, Scatter plots of E-P pair expression (triplicate averages) in plasmid vs integrated format for each of upstream, downstream, and STARR-seq enhancer positions. Boxes above scatter plots indicate the value of each correlation with darker shading indicating stronger correlation. **G**-**H**, Correlation of E-P pair expression (log2, triplicate averages, lower triangle) in the upstream, downstream, and STARR-seq assays’ enhancer positions in plasmid (**G**) and integrated (**H**) templates. Diagonals are scatter plots of biological replicates of each assay. “*R*” is Pearson’s correlation, and “n” is the number of E-P pairs in each scatter plot. The upper triangle is shaded according to the strength of the correlation, on the same scale as **D**, and labeled with the value of the correlation. See **Fig. S1** for the full correlation matrix with all assays. **F**, Each assay more closely matches a different proposed pattern of enhancer-promoter compatibility despite testing the same E-P pairs. A majority of variance in STARR-seq data is explained by intrinsic enhancer activity (x-axis) as in broad/equal compatibility while downstream data exhibits promoter-dependent scaling of enhancer effects as in the promoter responsiveness pattern. Upstream data exhibits the promoter responsiveness pattern but also exhibits apparent specificity in that the strongest enhancer activates the less responsive Promoter 1 (black) by a greater fold-change than the more responsive Promoter 2 (red). Promoter 2 corresponds to the genomic promoter sequence fragment from the endogenously highly expressed *FTH1* gene. Promoter 1 is the “noTFBS” sequence previously designed to be an inactive element^47^. See **Fig. S8** for all 6 assays. Note that the x-axis order of enhancers in each plot reflects the per-assay measurements and is not fixed, which we investigate in Fig. 2.

In considering possible sources of discrepancies in these observations, we noted that previous studies have used reporter assays with distinct designs. One variable is whether the reporter gene is encoded in an episomal vector or integrated into the genome, where the difference in chromatin context differs from endogenous loci and is suspected to influence transcriptional regulation^40–43^. A second variable is the placement of the enhancer either upstream, downstream, or inside of the reporter transcript (‘upstream’ MPRA ^35,39,44^, ‘downstream’ MPRA ^32,36,38^, and STARR-seq designs ^27,33,34^, respectively; **Fig. 1C**). While others have compared different MPRA assays and found discrepancies in their measurements of enhancer activities^42,43,45,46^, the ability of these assays to accurately measure enhancer-promoter compatibility have not been directly compared. Furthermore, none have been quantitatively compared with orthogonal genomic enhancer perturbations at corresponding native loci. These disagreements among MPRA-based studies, and the failure to connect them to orthogonal measurements at native loci, have left the field without a coherent framework for enhancer-promoter regulation.

In this study, we set out to establish a unifying framework for how intrinsic enhancer and promoter activities combine. We systematically assessed 6 reporter assay designs and identified technical confounders that distort apparent patterns of enhancer-promoter compatibility. We then developed and applied new reporter assays that avoid these confounders and closely mirror endogenous enhancer and promoter activities. Our results support and expand prior observations of differential promoter responsiveness: promoters differed dramatically in their responsiveness (1.1-fold to >100-fold activation in response to strong enhancers), while the rank-order of enhancer activation was similar across promoters. To demonstrate their utility beyond the reporter context, we directly incorporated the promoter responsiveness measurements from the MPRA into the ABC model, improving the quantitative prediction of endogenous enhancer-gene regulation measured by CRISPR perturbation experiments. Responsiveness itself was shaped by both classical core promoter motifs and other transcription factor motifs proximal to the transcription start site. Overall, this framework accounts for observations of both broad enhancer promiscuity and specific enhancer skipping and can explain how housekeeping genes maintain stable expression despite dynamic enhancer landscapes. Together, these findings establish that promoter responsiveness to transcriptional activation is an intrinsic, sequence-based property of promoters that strongly influences enhancer regulation in the human genome.

## Results

### Reporter assay design confounds measurements of E-P compatibility

To understand how reporter construct design or plasmid delivery of MPRAs might influence measurements of enhancer-promoter (E-P) compatibility, we tested 6 MPRA assays: 3 enhancer locations (upstream, downstream, or STARR-seq-style) delivered either episomally or integrated into chromatin (**Fig. 1C**). The 3 enhancer locations mirror reporter construct designs used in previous studies, changing only the position of the enhancer relative to the promoter and reporter gene while keeping the rest of the sequence identical: (i) “upstream” places the enhancer sequence proximally upstream of the promoter sequence and gene body (as in the commonly used “MPRA” ^48,49^ or “lentiMPRA” ^44,50^ assays), (ii) “downstream” places the enhancer sequence proximally downstream of the polyadenylation signal (similar to ^32,36^), and (iii) “STARR-seq” places the enhancer sequence within the transcribed gene body (as in ^27,33,34,51^). To also test whether E-P compatibility might differ between episomal and genomic contexts, each of the 3 constructs were cloned into the same plasmid backbone and delivered by either (i) transient transfection (*i.e.*, as for most previous MPRA studies ^52^) or (ii) stable integration into the genome at the *AAVS1* safe harbor locus ^47,53,54^.

To compare the 6 assays, we tested the same pool of 21 enhancer sequences x 26 promoter sequences (“21E x 26P”, 546 E-P pairs) in K562 cells. The enhancers and promoters were selected to include positive and negative control sequences with a large range of activities based on previous reporter assays ^33,47,50,53^. Hereafter, we refer to sequences cloned in the enhancer position as “enhancers” and sequences cloned in the promoter position as “promoters.” The regulatory function of these elements may differ from their naming, which we investigate below.

In brief, we cloned the 546 E-P pairs into each of the 6 assays and transfected 10 million K562 cells in 3 biological replicates each. We harvested pA-enriched RNA and genomic DNA either 24 hours later (for episomal assays) or 17 days later (7 days of puromycin selection and 10 days of post-selection recovery, for AAVS1-integrated assays). We then read out RNA expression via sequencing of barcodes unique to each E-P in separately extracted RNA and DNA. We computed reporter expression as the ratio of read-depth normalized RNA counts to DNA counts averaged over barcodes.

While reporter expression was highly technically reproducible within each assay (biological replicate Pearson’s *R* > 0.97, **Fig. S1A**), these measurements showed variable levels of concordance across assays (Pearson’s *R* = 0.67–0.93, **Fig. S1A**). The key variable was enhancer location, rather than plasmid vs in-genome. Plasmid and AAVS1-integrated assays with the same enhancer location (*e.g.*, upstream plasmid versus upstream integrated) were highly correlated (Pearson’s *R*=0.91-0.93) (**Fig. 1D**). Upstream versus downstream assays were strongly correlated with each other (*R*=0.91 and *R*=0.89 for integrated and plasmid assay comparisons, respectively) but were less well correlated with the STARR-seq configuration (*R*=0.75-0.83) (**Fig. 1E**). Thus, in our system, discrepancies in reporter expression were primarily driven by differences in enhancer location rather than differences between an episomal and *AAVS1*-integrated chromatin context.

To test if reporter construct design with respect to enhancer location could yield qualitatively different patterns of E-P compatibility, we evaluated the correspondence of each assay with the 3 models of E-P compatibility previously proposed: broad compatibility ^33,34^, combinatorial specificity^21–23,25,27,35^, and promoter responsiveness ^36–39^ (**Figs. 1B**). We first computed “basal promoter strength” as the average expression of a promoter when paired with negative control enhancers. We defined “enhancer effects” as the fold-change by which an enhancer activates a promoter over its basal activity. We defined “enhancer activity” (*E_avg_*) and as the effects of an enhancer averaged across all promoters.

The three assay designs indeed yielded different patterns of compatibility. In STARR-seq, enhancers had similar effects across promoters (*R*^2^=0.715), consistent with broad compatibility (**Fig. 1F**). In the downstream assay, enhancers had similar rank order of effects on promoters but with different magnitudes of effect, consistent with the promoter responsiveness pattern (**Fig. 1F**). In the upstream assay, the data appeared similar to the promoter responsiveness pattern, but certain pairs of enhancers and promoters appeared to show unusually strong activation, suggestive of some combinatorial specificity (**Fig. 1F, Fig. S8**). These assay design-specific biases were consistent across both episomal and *AAVS1*-integrated chromatin contexts (**Fig. S8**).

Thus, on the same set of enhancer and promoter sequences, the design of the reporter construct changes the interpretation of E-P compatibility patterns.

### Resolving technical confounders in reporter assays unifies patterns of E-P compatibility

To resolve these discrepancies, we examined positive and negative controls for promoter activity, enhancer activity, and E-P compatibility. The designations of positive and negative controls were critical to this assessment and described in **Methods**. Briefly, negative control promoter sequences include designed inactive elements and inaccessible chromatin regions. Positive control enhancers in the 21E library are all marked by DNase I hypersensitivity, H3K27ac ChIP-seq signal, and P300 ChIP-seq signal at their corresponding native loci and validated in other assays of enhancer activity. We observed that basal promoter strengths were highly correlated across all assays and could not explain discordant E-P pair expression (**Fig. S1D**). All assays cleanly separated the expression levels of promoters from highly expressed genes versus negative control promoters (**Fig. S1B**). Enhancer activities measurements were less consistent across assays (**Figs. 2A, S1C, S1E**). Investigating the discordant measurements of enhancer activity, we pinpointed two technical confounders of E-P compatibility measurements:

**Figure 2.**
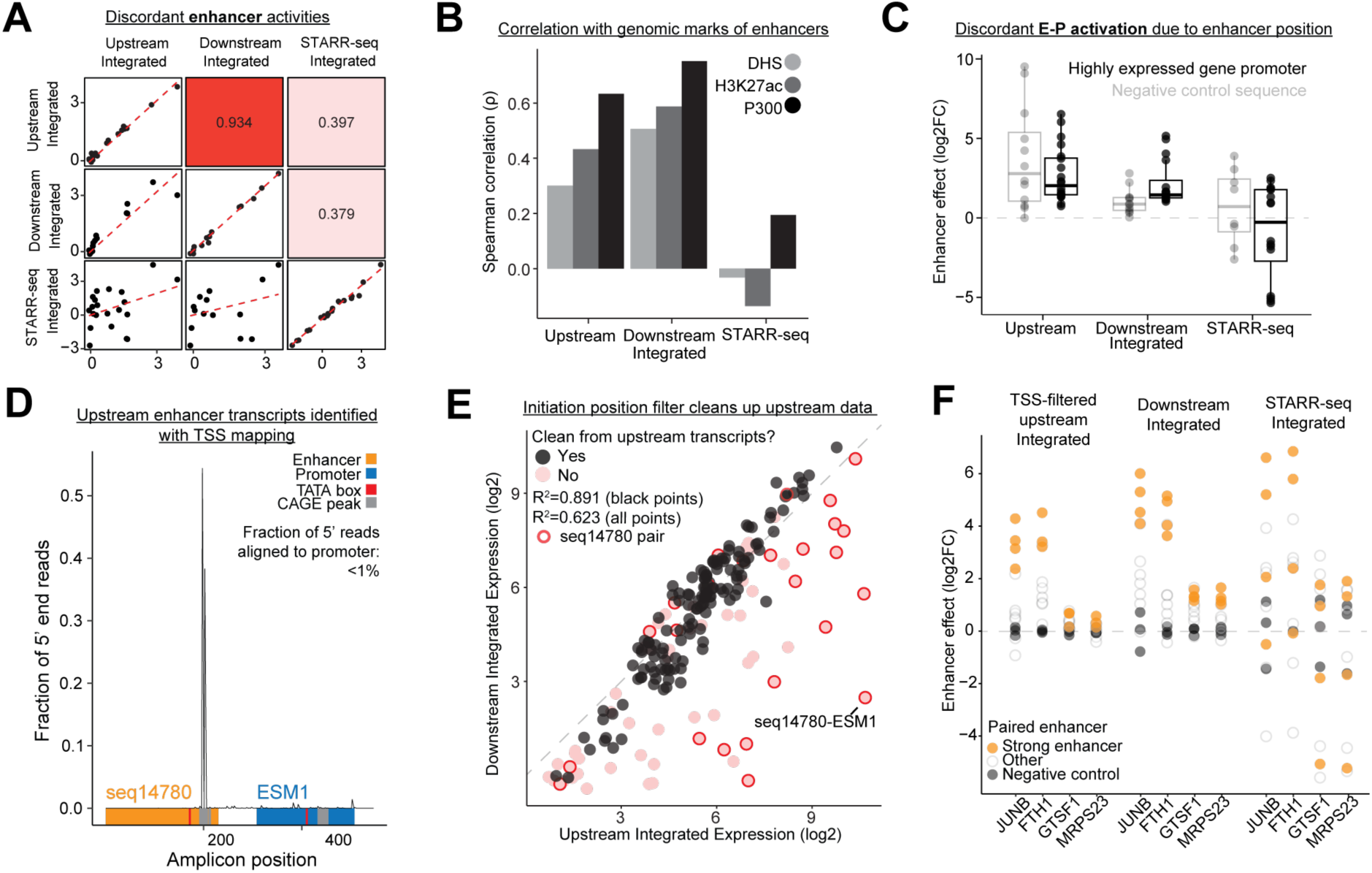
Assessing the validity of E-P compatibility measurements across assays. **A**, Scatter plots comparing the enhancer activities (*E_avg_*, triplicate averages) computed with respect to the *AAVS1*-integrated versions of each assay (lower triangle). Biological replicates are plotted on the diagonals. The upper triangle is shaded according to the magnitude of the Pearson’s correlation of the corresponding lower triangle scatter plot. See **Fig. S1E** for the full correlation matrix with all assays. **B**, Correlation (Spearman’s rho) of the enhancer activities from *AAVS1*-integrated assays with epigenomic measurements of the corresponding enhancers at their native loci (DNase I hypersensitivity, H3K27ac ChIP-seq, P300 ChIP-seq). See **Fig. S1F** for all assays. **C**, Box plots of enhancer effects (log2FC) for 4 positive control enhancers paired with 4 negative control promoter sequences or 5 highly expressed gene promoters. See **Fig. S1G** for all assays tested. **D**, Distribution of reads from the 5’ end mapping of transcription initiation sites on the *AAVS1*-integrated, upstream reporter for the E-P pair seq14780-ESM1. The red “TATA box” indicates a consensus TATAAA within the sequence. Gray “CAGE peak” indicates an annotated CAGE peak (FANTOM5) at the corresponding genomic locus. **E**, Scatter plot of E-P pair expression in the upstream integrated assay versus the downstream integrated assay. Each E-P pair is colored by whether the proportion of 5’-end mapping reads that align to the intended promoter region is greater than 90% (black) or not (red). Pairs with low coverage in the 5’-end mapping assay (**Fig. S2C**) were filtered out in comparison to Fig. 1E. E-P pairs with seq14780 (an element with pathologically strong autonomous promoter activity) used as the enhancer sequence are outlined in red. **F**, Enhancer effects on two gene promoters with the largest average enhancer effects (*JUNB*, *FTH1*) and two gene promoters with the smallest average enhancer effects (*GTSF1*, *MRPS23*) in each *AAVS1*-integrated assay. “Strong enhancers” are positive controls (same as **Fig. S1C**) and “weak enhancers” are those without either positive or negative control designations.

STARR-seq measurements of enhancer activity appeared to be confounded by inclusion of the enhancer sequence in the RNA reporter transcript. Enhancer activities did not correlate well with the level of activating marks on the corresponding sequences in the genome (DNase I hypersensitivity and ChIP-seq for H3K27ac or P300) (**Figs. 2B, S1F**), showed poor agreement when cloned in forward versus reverse sequence orientations^55^ (**Fig. S3A-B**), and some CRISPR-validated strong enhancers even led to negative effects on reporter expression (**Fig. S1C**). Because enhancers in the STARR-seq assay are transcribed as part of the reporter RNA, we hypothesize that the tested sequences may not only have effects as DNA enhancers but also affect stability or processing of the RNA transcript ^55,56^. Consistent with this hypothesis, long-read RNA sequencing revealed several alternatively spliced reporter transcripts using cryptic splice acceptors and donors rather than the designated acceptor and donor, similar to a previous study finding cryptic splicing as a confounder in 3’ UTR MPRAs ^57^ (**Fig. S2D**). Curiously, effects on RNA stability or processing may not be the only explanation for this discordance. Enhancers with similar strengths in both forward and reverse orientations in STARR-seq, which might be expected to control for RNA-specific effects by changing RNA sequence, still significantly deviated from their measured activity in an upstream position (**Fig. S3C**). Ultimately, these and other confounders in STARR-seq inflate the fraction of variance in reporter expression attributable to apparent enhancer activity, leading to an interpretation of these data as more consistent with broad/equal E-P compatibility (**Figs. 2C, S3D-F, 2F**).

Upstream MPRA measurements appear to be confounded by the ability of some enhancer sequences to act similarly to promoters and autonomously initiate transcription. This was most apparent in cases where enhancers strongly activated negative control sequences (**Figs. 2C, S1G**). We developed an approach similar to RAMPAGE and STRT-seq ^58,59^ to directly map the 5’ ends of the reporter transcripts from each E-P pair (**S2A, S2B**). For approximately half of measured E-P pairs, a significant portion of reporter transcripts initiated in the enhancer sequence, rather than in the promoter sequence (**Figs. 2D, S2C**). Basally weak promoters, such as ESM1, minCMV, and minP, tended to be dominated by enhancer-initiated transcripts regardless of enhancer strength (**Fig. 2D, S2C**). Basally strong promoter sequences were largely unaffected, except when paired with enhancer sequences with very strong autonomous promoter activity (e.g. “seq14780”, the most active sequence from a K562 genome-wide, upstream lentiMPRA screen^50^) (**Fig. 2E, S2C**). Notably, strong autonomous promoter activity and strong enhancer activity were separable properties. Enhancer sequences “seq14780” and “chr3:128134842-128135106-enhancer_rc” were similarly strong enhancers in downstream MPRAs yet only the former had strong, autonomous promoter activity (**Table S6**, **Figs. S1C**, **S2G**). Enhancer-initiated transcription also did not simply increase expression at all paired promoters uniformly. For example, the most highly expressed enhancer promoter pair in the integrated upstream assay was seq14780 paired with ESM1, a basally weak promoter, rather than any of the basally more active promoters (**Fig. 2E**). This suggests that there are non-additive effects from placing promoters in proximity that remain to be explored. Lastly, we also observed cases with full transcript, long-read sequencing where enhancer-initiated transcripts could use cryptic splice sites rather than those of the designed intron, altering transcript stability and further confounding barcode-only measurements (**Fig. S2D**). Overall, capturing full-length reporter transcripts with our 5’-end mapping assay revealed several unpredictable confounders of expression in upstream assays resulting from enhancer-initiated transcription.

To resolve confounding by enhancer-initiated transcripts in upstream MPRAs, we excluded enhancers with strong autonomous promoter activity and promoters with weak autonomous promoter activity (“TSS-filtered upstream MPRA”). We found that all E-P pairs more highly expressed in the unfiltered upstream MPRA compared to the downstream assay were confounded by enhancer-initiated transcripts (**Fig. 2E**). After filtering out these pairs, the TSS-filtered upstream MPRA and downstream MPRA yielded consistent patterns of enhancer-promoter compatibility (**Figs. 2F**, **S2E**, **S8**). Thus, enhancer-initiated transcripts in the upstream MPRA assay lead to the confounded appearance of combinatorial E-P specificity.

Together, these observations identify and reconcile discrepancies in E-P compatibility measurements that result from assay design. Downstream MPRA or TSS-filtered upstream MPRA assays provide concordant measurements of E-P compatibility, avoiding confounding by effects of 1) enhancer sequences being placed within the transcript or 2) enhancer-initiated transcription.

### Reassessing E-P compatibility reveals a large dynamic range in promoter responsiveness

To more broadly test how enhancer and promoter activities combine while avoiding the confounders we previously identified, we designed a larger MPRA library with 203 enhancer and 78 promoter sequences (“203E x 78P”) selected from 7 genomic loci in which we and others have conducted CRISPR screens to comprehensively map which enhancers regulate which genes ^1,5,60^ (**Fig. 3A)**. We conducted the TSS-filtered upstream MPRA assay integrated at *AAVS1* and, by applying 5’-end readouts, successfully filtered out enhancer-initiated transcription (**Figs. S2F, S2G**). We obtained high replicate reproducibility and concordance with endogenous promoter and enhancer activities (**Figs. S4A-G**).

**Figure 3.**
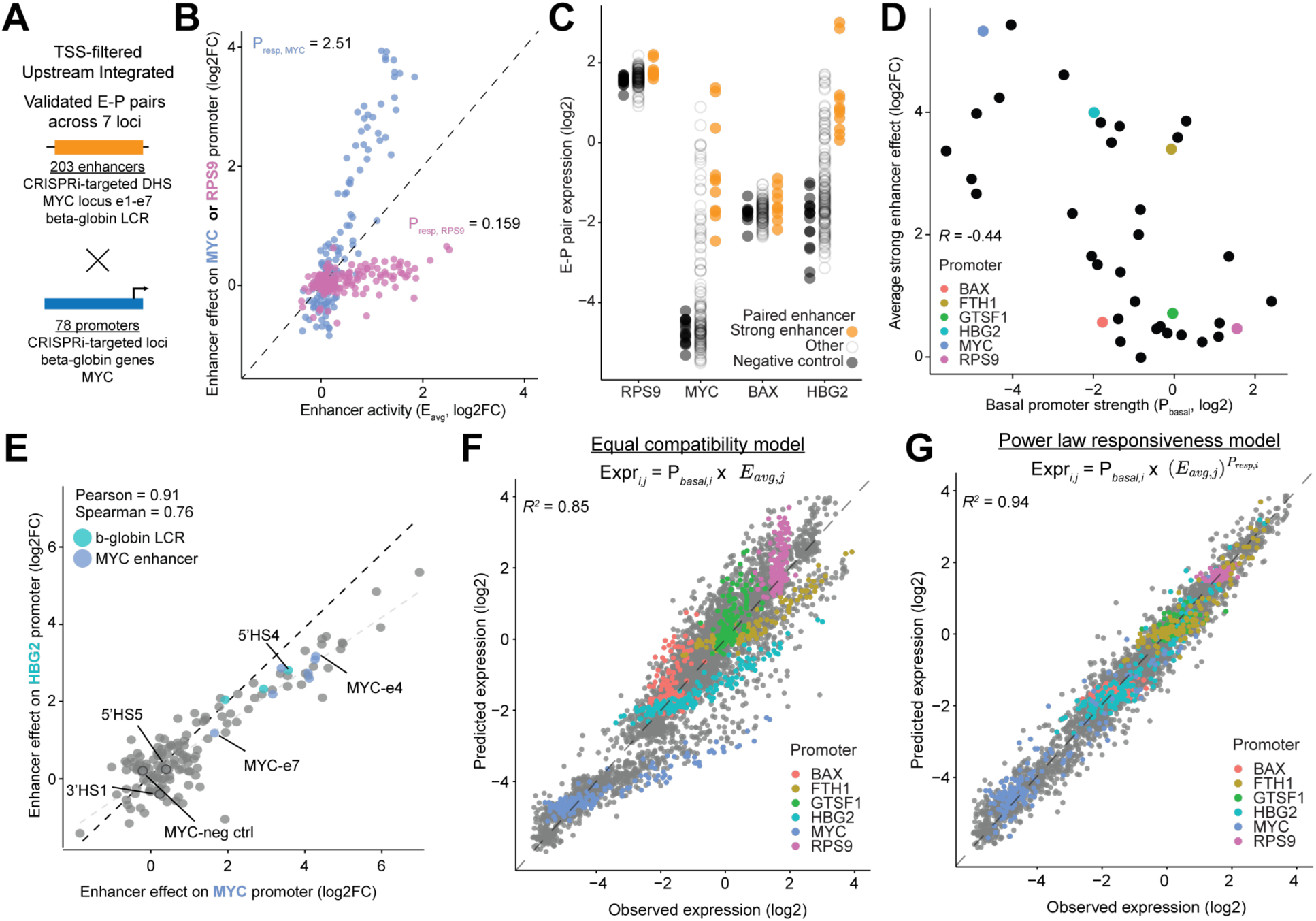
Promoters differentially scale the effects of enhancer activation. **A**, Library of 203 enhancer sequences and 78 promoter sequences tiling well-validated E-P pairs from 7 genomic loci. **B**, Scatter plot of enhancer effects on the *MYC* and *RPS9* promoters versus the enhancer activity (*E_avg_*) over all promoters, for each enhancer sequence (dots). **C**, E-P pair expression of 4 promoters corresponding to *RPS9*, *MYC*, *BAX*, and *HBG2* with 3 classes of enhancer sequence. Black dots indicate pairs where the enhancer is a negative control used to compute basal expression (*P_basal_*). Orange dots indicate pairs where the enhancer is an expected strong enhancer used to compute the average strong enhancer effect in the next panel. Gray dots are all other pairs. **D**, Average enhancer effect of selected strong enhancers on each promoter (log_2_FC) versus the promoter’s basal strength (*P_basal_*). Only promoter sequences passing the 5’-end mapping TSS filter are plotted. **E**, Enhancer effect of each enhancer sequence on *HBG2* (y-axis) and *MYC* (x-axis), respectively. Tiles of well-validated enhancers and negative controls from each respective locus are highlighted. Black dashed line represents y = x, and the gray dashed line is the line of best fit. **F**, Predicted versus observed log expression of all E-P pairs (dots) in the library under a simple multiplicative model of basal promoter strength multiplied by enhancer effect. A selection of promoters that are more and less responsive to enhancer sequences are highlighted. **G**, Same as **F**, with the same promoters highlighted except the equal compatibility model is modified with an exponential factor adjustment to enhancer effects representing the responsiveness of the promoter.

In concordance with our previous, smaller E-P library, promoters showed substantial differences in their responsiveness to enhancers. For example, the strongest enhancers activated the promoter of *MYC* by 30-fold, and the promoter of *RPS9* (a “housekeeping” gene) by <2-fold (**Fig. 3B**). These differences were not well explained by “saturation” of transcription at the high expression levels: basally weak promoters with high responsiveness, when paired with a strong enhancer, could achieve higher expression than basally strong promoters with low responsiveness (**Fig. 3C**). For example, the *BAX* and *HBG2* promoters exhibited similar basal promoter activity but were activated <2-fold and 16-fold by strong enhancers, respectively (**Fig. 3C**). Promoter responsiveness was only moderately correlated with basal promoter strength (Pearson’s R = -0.44, **Fig. 3D**). These data suggest promoters exhibit two independent, intrinsic characteristics: basal activity and responsiveness to enhancer inputs.

Across the library of validated and predicted enhancers, we observed consistent rank-ordering of enhancer activity enhancers at different promoters; however, the absolute magnitudes of these effects scaled with promoter responsiveness. For example, enhancer sequences activated the responsive *MYC* and *HBG2* promoters with similar rank order, though with different dynamic ranges (Pearson’s *R* = 0.91, Spearman *rho* = 0.76, **Figs. 3E, S5A**). Across promoters with average strong enhancer activation above 4-fold, enhancer effects were highly correlated (average Pearson’s *R* = 0.89 and Spearman *rho* = 0.74) **(Fig S5A, S5B)**. Thus, promoters responded to enhancers with similar rank-orders but with different slopes in log-log space.

In order to capture the promoter-specific scaling of enhancer activation, we attempted to model promoter expression across E-P pairs in two ways: (1) a model that assumes equal, multiplicative effects of all enhancers on all promoters, and (2) a model in which enhancer activity is raised to the power of a promoter-dependent, responsiveness parameter (*P_resp_*).

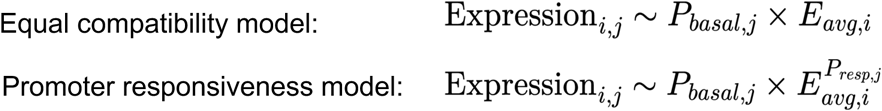

While the “equal compatibility” model explained some of the variance in the expression across constructs (*R*^2^=0.85), promoters clearly had distinct slopes in log-log space (**Fig. 3F)**. The promoter responsiveness model explained 94% of the variance in the data — very close to the correspondence between biological replicates (*R*^2^=0.96, **Fig. 3G, S5D**). Together, this indicates that enhancers are broadly compatible but that promoters differ dramatically in their responsiveness. The relationship between expression and enhancer activity appears to follow a power law, with different promoters having different exponents.

### Promoter responsiveness predicts enhancer effects in genome

We next tested whether our promoter responsiveness model predicted E-P regulatory interactions not only in a reporter assay but at their endogenous locations in the genome. We began by leveraging the Activity-by-Contact (ABC) model, which encodes the notion that enhancers activate nearby promoters proportional to their intrinsic enhancer activity multiplied by the frequency of E-P contact in 3D space^1^. Empirically, ABC has been found to predict which enhancers have regulatory effects on which nearby genes in CRISPR perturbation experiments significantly better than distance-to-TSS baselines (area under the precision-recall curve (AUPRC) = 0.65 vs 0.39). Yet, the ABC model has not been able to explain certain complexities in the CRISPR data, such as why enhancers sometimes “skip” over a nearby gene to regulate a more distant one and why ubiquitously expressed genes are insensitive to enhancer perturbations^1,5^ (59% precision at 70% recall). ABC also does not accurately predict enhancer effect sizes. We hypothesized that integrating promoter responsiveness as an additional factor could improve the ability of ABC to explain E-P regulation in the genome ^13^.

We first compared our 203E x 78P library MPRA measurements to the CRISPR data of matched enhancers and promoters in the genome. In this data originally used to benchmark the ABC model, CRISPRi was used to perturb every DNase I hypersensitive (DHS) element within 450 kb of 30 selected genes across 5 genomic regions. Responsiveness (*P_resp_*) was positively correlated with the number of CRISPR-validated enhancers for the corresponding gene in the genome (Pearson’s *R* = 0.80, *P* = 7.86 × 10^−9^)—promoters with high *P_resp_* had up to 4 enhancers, whereas promoters with the lowest *P_resp_* often had none (**Figs. 4A, 4B**). *P_resp_* was also correlated with enhancer effect sizes (*R*=-0.64, *P*=1.17 × 10^−6^) (**Fig. 4C**). Among promoters with low responsiveness (*P_resp_* < 1), the strongest enhancer had just a 25% effect on gene expression, while among the responsive promoters (*P_resp_* > 1), multiple genes had enhancers with >50% effects.

**Figure 4.**
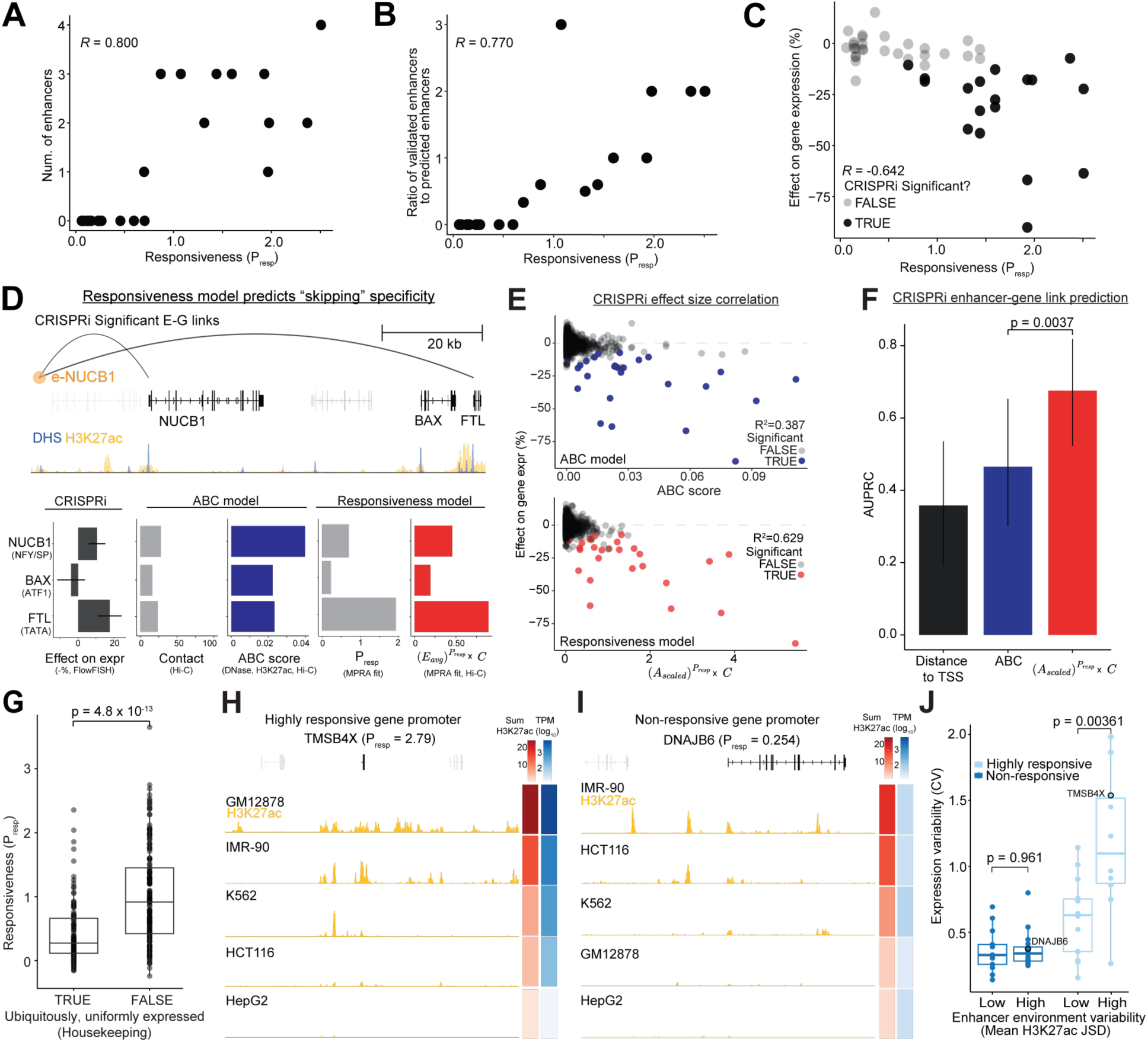
Promoter responsiveness tunes sensitivity to *cis*-regulatory signals at endogenous loci. **A**, The number of CRISPR-validated enhancers per gene at previously studied loci versus MPRA-measured, gene promoter responsiveness. **B**, Ratio of the number of CRISPR-validated enhancers to the number of ABC-predicted enhancers (ABC score > 0.02) per gene versus gene promoter responsiveness. A ratio greater than 1 indicates that there were more CRISPRi-validated enhancers than there were enhancers predicted by ABC. **C**, Genes with more responsive promoters are more strongly affected by ABC-predicted enhancers. Effect of strong putative enhancers (ABC score > 0.02) from CRISPR experiments on target genes plotted against the target gene promoter’s responsiveness. **D**, An intergenic enhancer (“e-NUCB1”, orange circle) characterized via CRISPRi regulates *NUCB1* and *FTL* but “skips” *BAX* (hg38 coordinates: chr19:48877370-48967448). Corresponding DNase I hypersensitivity and H3K27ac ChIP-seq signal tracks for the locus are shown below. Bar plots indicate 3D contact, Activity-by-Contact (ABC) score, MPRA-measured responsiveness of the three gene promoters, and responsiveness incorporated into the R-ABC model using MPRA-measured enhancer activity and promoter responsiveness. **E**, Incorporating responsiveness into ABC scores (bottom) improves correlation with measured CRISPRi enhancer perturbation effect sizes compared to ABC alone (top) across a tiling screen of >1,600 element-gene pairs across 6 loci (25 significant pairs, 1627 non-significant pairs). Statistically significant effect sizes from the CRISPRi screen are colored in red (bottom) or blue (top) while insignificant ones are colored in gray. **F**, Classification performance of enhancer-gene pairs from the same CRISPRi data improves upon incorporating responsiveness (red) compared to ABC (blue) and a baseline TSS distance predictor (black). *P-*value is obtained by testing the difference in AUPRC between the Responsiveness and ABC models using n=10,000 bootstrap samples. **G,** Comparison of *P_resp_* measurements for promoters of genes labeled as ubiquitously and uniformly expressed or not based on previous analysis^13^ of 1,829 CAGE experiments from the FANTOM5 consortium^61^. **H**,**I**, Increasing normalized H3K27ac ChIP-seq signal across 5 cell lines corresponds to increased RNA-seq expression and expression variability for a highly responsive gene promoter *TMSB4X* (**H**) but not a non-responsive gene promoter *DNAJB6* (**I**). **J**, Expression variability of a given gene (coefficient of variation (CV) computed over cell lines with measured RNA-seq TPM>1) across the 5 cell lines significantly increases with H3K27ac ChIP-seq signal profile variability for genes with highly responsive promoters (top quartile) while it does not change for genes with non-responsive promoters (bottom quartile). High and low enhancer environment variability corresponds to the top and bottom quartiles of all genes of mean, pairwise H3K27ac ChIP-seq profile Jensen-Shannon distance using a 200 kb TSS-centered window per gene.

To predict the quantitative effects of enhancers on gene expression, we incorporated *P_resp_* into the ABC model as an exponent of enhancer activity, as in Eq. 2, to formulate the Responsiveness-incorporated ABC (R-ABC) model:

ABC model numerator:

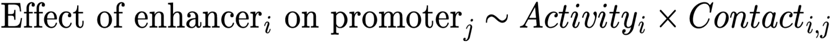

R-ABC model:

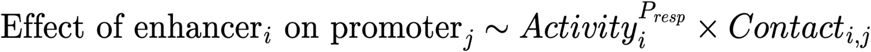

We illustrate the calculation with an example locus in **Fig. 4D**, where we previously measured the effect of perturbing an enhancer on 3 nearby genes: *NUCB1*, *BAX,* and *FTL*. CRISPRi-mediated inhibition of the enhancer significantly decreased the expression of *NUCB1* and *FTL* expression (by 11% and 18%, respectively) but had no significant effect on *BAX* expression, suggesting that the enhancer “skips” *BAX* to regulate the more distal *FTL*. The ABC model does not explain this behavior: all 3 genes show similar levels of 3D contact with the enhancer, and the ABC scores predict that the enhancer should regulate all 3 promoters with *NUCB1* as the most strongly regulated (**Fig. 4D**). Our MPRA measurements, however, showed a hierarchy of promoter responsiveness: *FTL* (*P_resp_*=1.93) is much more responsive than *NUCB1* (*P_resp_*=0.700), and both are more responsive than *BAX* (*P_resp_*=0.229). By incorporating *P_resp_* into the ABC score, the R-ABC model more accurately reflected the enhancer’s relative effects on the 3 genes, correctly predicting the strongest regulation for *FTL* and the apparent “skipping” behavior for *BAX*. This example shows how the improved R-ABC model captures an example of promoter-skipping not captured by the ABC model, which assumes equal compatibility between all enhancers and promoters.

To extend this analysis at all CRISPR-validated loci, we computed ABC scores and R-ABC scores for 1,606 distal element-gene pairs, across 6 loci, for which we had both CRISPRi and MPRA measurements. The R-ABC model explained 63% of the variance in the CRISPR-measured effect sizes, a significant increase over the ABC model (39%) (**Fig. 4E**). Similarly, R-ABC achieved better binary classification performance than ABC (AUPRC = 0.676 vs 0.466, *P* = 0.0037 (bootstrap)) (**Fig. 4F**).

Thus, our measurements of promoter responsiveness and the power law equation derived from E-P MPRAs generalizes to distal regulation in the genome. The R-ABC model incorporates promoter scaling of enhancer activity and improves upon the ABC model for predicting enhancer-gene regulation, explaining certain cases where enhancers appear to “skip” over nearby genes.

### Promoter responsiveness tunes sensitivity to cis-regulatory input across cell types

To study the role of promoter responsiveness in endogenous regulation with more promoters than those in the matched CRISPR dataset, we expanded our measurements of *P_resp_* with an additional 500 promoter library. We conducted another TSS-filtered upstream MPRA assay integrated at *AAVS1* with 500 promoters chosen from a range of endogenous gene expression levels, each combined with the original 21E enhancer library (“21E x 500P” library) (**Fig. 5A**). As with the 203E x 78P library, E-P pair expression was well modeled by the Promoter Responsiveness model (Equation 2) (*R*^2^ = 0.98, **Figs. S5C, S5G**) and estimates of *P_resp_* were consistent for promoters shared between the two libraries (Pearson’s *R* = 0.94, **Fig. S5H**).

**Figure 5.**
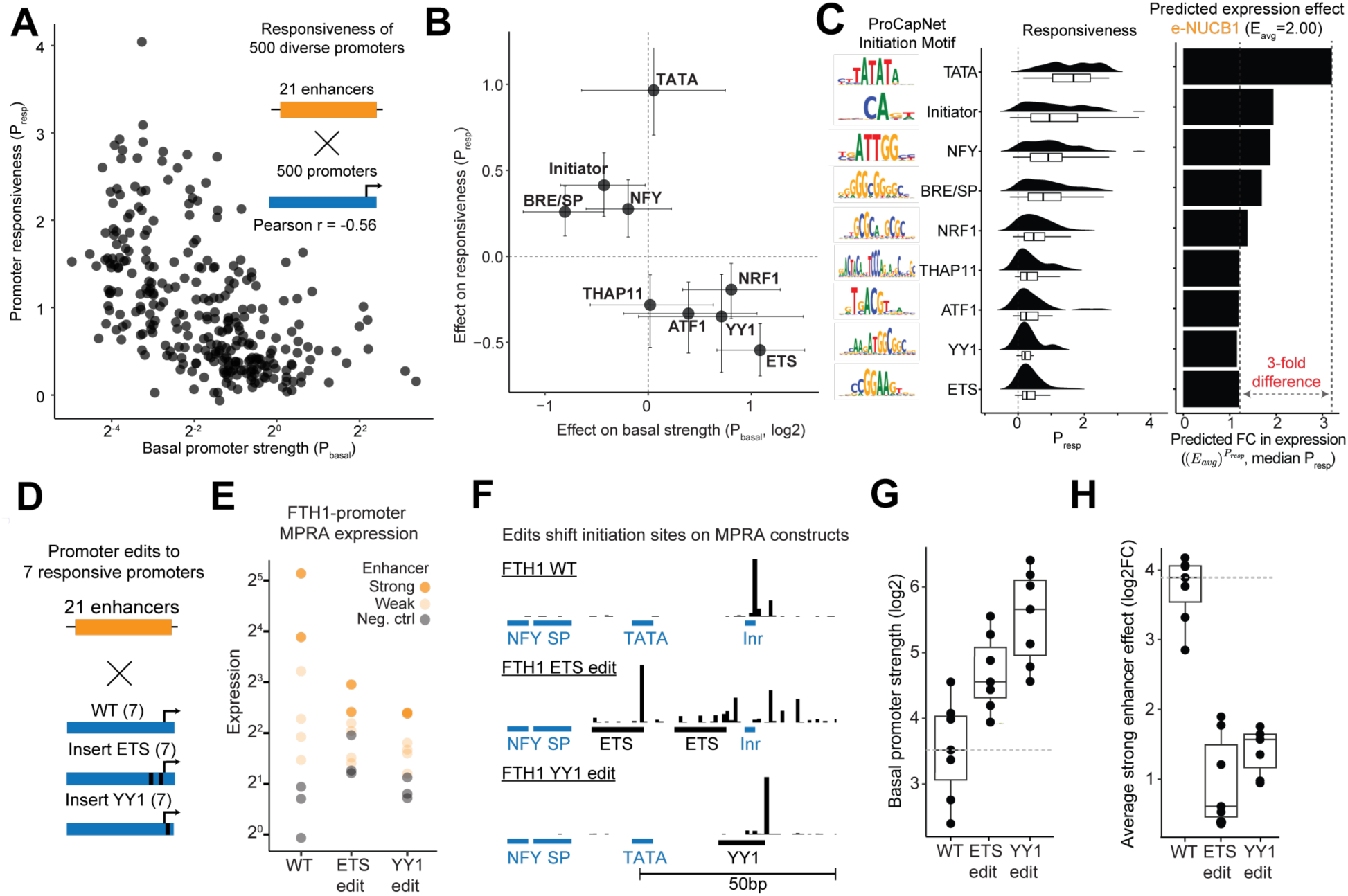
Sequence determinants of promoter responsiveness. **A**, Scatter plot of promoter basal activity (*P_basal_*) and responsiveness (*P_resp_*) with a library of 21 enhancers by 500 promoters. **B**, Effect of the presence versus absence of ProCapNet motifs on *P_basal_* (log_2_) and *P_resp_*. Effects and 95% confidence intervals were estimated by Wilcoxon test (Hodges-Lehmann median estimates). See **Fig. S6C** for plots of **A** faceted by the motifs plotted here. **C**, (left) Histograms and box plots of *P_resp_* values for promoters containing each ProCapNet motif. (right) Predicted effect of motif presence on activation by an example enhancer, e-*NUCB1*, calculated by raising enhancer activity (*E_avg_* = 2.00) to the power of the median responsiveness of promoters containing each motif. **D**, Promoter motif edit library consists of the same 21 enhancers with seven sets of wild-type and edited promoters (ETS inserted at -30 and -10, and YY1 inserted at +1 relative to the RefSeq annotated TSS). The motif insertions replace the original sequence at the positions indicated. **E**, Expression of each E-P pair for FTH1 wild-type and edited promoters using the same enhancer classifications as Fig. 2F. **F**, Promoter motif annotations and 5’ end mapped initiation sites on the wild-type *FTH1* promoter sequence and two edited *FTH1* promoter sequences in the MPRA construct. **G**,**H**, Basal promoter strength (*P_basal_*) and enhancer effect averaged over 4 strong enhancers for the seven sets of wild-type and edited promoters.

We hypothesized that promoter responsiveness could help explain why ubiquitously and uniformly expressed genes (housekeeping genes^62^) are 5-fold less likely to have distal regulatory elements compared to cell-type–specific genes^1,13^. Consistent with this hypothesis, genes previously annotated as ubiquitously and uniformly expressed^13^ based on 1,829 CAGE experiments from the FANTOM5 Consortium^61^ had significantly lower *P_resp_* than cell-type–specific genes (average *P_resp_* = 0.43 vs 1.0, respectively, Wilcoxon rank-sum test *p-*value = 4.8 × 10^−13^) (**Fig. 4G**). This suggests that ubiquitously and uniformly expressed genes typically have lower *P_resp_*, which may act to insulate gene expression from differences in enhancer activity across cell types, not just in K562 cells. In contrast, cell-type–specific genes with high *P_resp_* may amplify those differences into greater ranges in gene expression.

To test whether variation in *cis*-regulatory input produces different ranges of expression depending on the promoter’s responsiveness, we compared variation in gene expression with variation in active chromatin marks across 5 ENCODE cell lines for genes with high and low *P_resp_*. Operationally, we estimated variation in gene expression with RNA-seq data (coefficient of variation) and variation in *cis*-regulatory inputs with the variance in H3K27ac ChIP-seq profiles similarity (mean Jensen-Shannon distance (JSD)) in a 200-Kb window surrounding the promoter which is likely to contain most regulatory enhancers. For example, *TMSB4X* (*P_resp_* = 2.8) and *DNAJB6* (*P_resp_* = 0.25) exhibited similar variability in their cis-regulatory environments (mean JSD across cell types = 0.831 vs 0.832) but have *P_resp_* values in the top and bottom quartiles, respectively. In the genome, *TMSB4X* expression across cell types spans three orders of magnitude and correlates with the sum of nearby H3K27ac signals, while *DNAJB6* is more uniformly expressed despite changes in its nearby H3K27ac signals (**Figs. 4H, 4I**). Across all genes with corresponding *P_resp_* measurements, those in the bottom quartile of *P*_resp_ showed low variation in expression across cell types regardless of the level of variation in nearby *cis*-regulatory input (**Fig. 4J**). In contrast, top-quartile *P_resp_* promoters showed approximately 2-fold greater variation in gene expression when *cis*-regulatory input variation was high (top quartile) versus low (bottom quartile). Thus, promoter responsiveness tunes how sensitively target gene expression responds to variation in *cis*-regulatory input across cell types and helps to explain how housekeeping genes with low *P_resp_* maintain stable expression despite changes in their regulatory environment.

### Sequence determinants of promoter responsiveness

To understand how promoter responsiveness might be encoded in DNA sequence, we examined whether core promoter motifs and transcription factor motifs correlated with responsiveness. While promoters are typically classified by classical core promoter motifs such as TATA box, initiator, and DPE elements, these motifs occur at a minority of human promoters^63^. Thus, to conduct a more inclusive analysis, we leveraged ProCapNet, a deep learning model trained to predict genome-wide patterns of transcription initiation as measured by PRO-cap^64^ from DNA sequence.

We interrogated ProCapNet models previously trained on K562 PRO-cap data^64^ to assess the importance of 12 recurrent and high-scoring sequence features in our set of promoter sequences for which we had measured responsiveness and basal strengths (**Figs. S6A, S6B**). We first annotated each promoter by whether it had a significant contribution from one or more of the ProCapNet-called sequence motifs. We then assessed the association of each sequence motif with both *P_resp_* and *P_basal_*. We found 10 motifs that significantly correlated with responsiveness and four that significantly correlated with basal strength (**Fig. 5B-C, Fig. S6C**). Promoters predicted by ProCapNet to utilize ETS and YY1 motifs were among those with less responsive promoters, consistent with previous observations ^33^. Other sequence motifs associated with reduced responsiveness in tested promoters include those of THAP11, ATF1, and NRF1. Meanwhile, promoters predicted to utilize TATA, Initiator, NFY, and BRE/SP motifs were significantly more responsive than the average promoter we tested. Notably, the motifs correlated with *P_resp_* did not necessarily also correlate with *P_basal_*. While ETS and YY1 motifs were correlated with an increase in *P_basal_* and a decrease in *P_resp_*, the TATA motif was correlated with *P_resp_* but not with *P_basal_*. This result further suggests that responsiveness and basal strength may be independently tunable promoter properties, at least for TATA promoters.

We analyzed the promoters at the locus in **Fig. 4A** and found that the two genes that were significantly downregulated following CRISPRi enhancer perturbation, *FTL* and *NUCB1*, are predicted to utilize a TATA motif and an NFY and SP motif, respectively. The dominant initiation motif at the promoter for the “skipped” *BAX* gene, on the other hand, is dominated by an ATF1 motif associated with reduced responsiveness. These observations align with broader trends shown in **Fig. 5C** where TATA promoters were on average the most responsive while ATF1, YY1, and ETS promoters were the least. For the example enhancer in **Fig. 4D**, this difference in responsiveness predicts a 3-fold difference in enhancer-driven expression—consistent with the observed differences among measured target genes.

We further compared *P_resp_* with the occupancy of transcription factors and cofactors at corresponding genomic promoters using ENCODE ChIP-seq data (**Fig. S6E**). We found several transcription factors correlated with reduced responsiveness including YY1 and ETS family transcription factors (GABPA, ELF1, ELK1, and ETV1) in concordance with ProCapNet-derived sequence features. The several cofactors correlated with reduced responsiveness included HCFC1, RBBP5, and PHF8, which have previously been shown to specifically act as co-activators for CpG-island promoters^16^. HCFC1 in particular has been shown to co-localize with multiple transcription factors associated with low *P_resp_* (ZNF143, THAP11, GABPA (an ETS family factor), and YY1)^65^. The concordance between sequence-based (ProCapNet) and occupancy-based (ChIP-seq) analyses, establish core promoter sequence identity as a primary determinant of responsiveness.

### ETS and YY1 motifs influence responsiveness and transcription start site selection

We next directly tested via sequence mutagenesis if two motifs enriched in low-*P_resp_* promoters, ETS and YY1, modulate responsiveness, and whether they do so by simply saturating transcriptional output at the endogenous TSS. We selected seven highly responsive promoter sequences and replaced the wild-type promoter sequence with ETS or YY1 motifs at specific positions relative to the TSS where they most frequently occur in our promoter sequences (-10 and -30 for ETS and +1 for YY1, **Figs. 5D, Fig. S6F**). In agreement with the enrichment analysis in **Fig. 5B**, directly inserting these motifs increased basal promoter strength and decreased responsiveness (**Figs. 5E, 5G, 5H**). Notably, we found that the decrease in *P_resp_* outweighed the increase in *P_basal_*, such that addition of these motifs effectively “stabilized” expression: edited promoters showed higher basal expression but lower maximum activated expression than wild-type. This result is inconsistent with the motifs saturating a rate-limiting step in transcription, which would result in increases in both basal and maximum, enhancer-activated expression levels. The expression levels of edited promoters confirms that TF motifs in the core promoter alter promoter activity and responsiveness.

Because both ETS and YY1 factors can directly contact general transcription factors and influence the pattern of transcription initiation, we assessed whether the motif edits modulated responsiveness through the original TSS or newly created initiation sites. We mapped the transcription start sites at the edited promoters and found that the ETS and YY1 motif edits altered the position and distribution of transcription start sites in all 7 promoters (**Figs. 5F**, **Fig. S7A**). For example, the original *FTH1* promoter sequence showed a focused transcription start site centered on an Initiator motif with a TATA box at –30 bp (**Fig. 5F**). Inserting two ETS motifs at -10 and -30 positions relative to the wild-type TSS led to a dispersed pattern of initiation directly over the inserted ETS motifs, and a near complete loss of initiation signal over the Initiator motif. Inserting a YY1 motif at the +1 position similarly led to a near complete loss of the original initiation site but created a single, focused TSS shifted 4-bp from the original TSS. The redistributed TSSs over the inserted motif in both edits broadly match the distribution initiation pattern at ETS motifs and focused initiation pattern at YY1 motifs learned by ProCapNet ^64^. Altogether, the observed shifts in TSS profiles suggest that the inserted motifs directly facilitate transcription initiation at the edited sites rather than cooperatively regulating transcription from the existing initiation site.

Together, these results demonstrate how introducing ETS and YY1 motifs in the core promoter can facilitate transcription initiation at new sites with reduced responsiveness to enhancer activation, rather than modulating activity at the endogenous TSS.

## Discussion

This study identifies transcriptional responsiveness as an DNA sequence-encoded property of core promoters that nonlinearly scales transcriptional activation by enhancers. Genes with the most responsive promoters are broadly activated by enhancers, while genes with the least responsive promoters are effectively insensitive to all enhancers. Previous studies using small numbers of enhancer-promoter pairs may have been selectively biased for responsive promoters suggesting broad promiscuity^32^ or focused on a single locus with a mix of responsive and non-responsive promoters that suggest promoter selectivity^22^. Meanwhile, high throughput MPRA-based studies have drawn divergent conclusions due to assay-specific confounders and different definitions of specificity^33–36^. Our view resolves paradoxical observations of both promoter selectivity and broad promiscuity of enhancers.

Promoter responsiveness scales activation by all enhancers and dramatically differs even among promoters of similar basal strength. Previous studies have observed variable promoter activation^23,39,66,67^. However, their reliance on just a few enhancers limited conclusions about whether this reflects categorical promoter preferences or a uniform scaling of all enhancer activation. Martinez-Ara et al. ^36^ proposed that promoters exponentiate enhancer activation across all enhancers according to a power law based on data from 8 promoters and 80 enhancers. However, differences in activation were entirely explained by basal promoter strength (*R* = –0.97). In this study, we show that responsiveness can significantly vary even among promoters with similar basal strength at a larger scale with 500 promoters and 200 enhancers (*R =* –0.44) (**Fig. 3C, 3D, 5A**). We also demonstrate that responsiveness has a greater dynamic range than previously observed: the most responsive promoter (from *GLUL*) is activated >100-fold, whereas the least responsive promoter (from *NUB1*) is not significantly activated by any enhancer. Thus, at loci where responsive and non-responsive promoters are interleaved, an enhancer might appear to be specific and “skip” over a nearby, non-responsive promoter to regulate a more distant, responsive promoter (*e.g.*, **Fig. 4D**). The magnitude of this effect establishes quantitative promoter responsiveness as a critical factor, along with 3D contact and enhancer activity, in predicting enhancer-gene regulatory interactions in the human genome.

What molecular mechanisms mediate promoter responsiveness? Our data provide several clues. First, we find that responsiveness is affected by DNA sequences including classical core promoter motifs such as TATA boxes at highly responsive promoters and binding sites for TFs like YY1 or ETS factors at non-responsive promoters. ETS and YY1 sites do not appear to be simply promoter-proximal enhancer elements — rather, they are enriched within 30bp of the TSS (**Fig. S6F**), bind factors that directly interact with core subunits of the TFIID complex^68,69^ , and can reposition the transcription start site ^70^ (**Figs. 5F, S7A**). Furthermore, while other promoter motifs like the TFIIB Recognition Element motif (BRE/SP) were broadly present in promoters across responsiveness classes, TATA boxes at promoters rarely co-occurred with ETS and YY1 motifs^71^ (**Fig. S6D**). Second, we find that the relationship between enhancer activity and promoter output follows a power law, with the exponent varying across promoters (**Fig. 3G**). Such a power-law relationship could arise, for example, from cooperative assembly of the transcriptional machinery or from a kinetic proofreading model, in which sequential rate-limiting steps must be traversed before productive elongation, both in subsaturating model regimes^72–74^. Apparently, the strong enhancers we tested here were still not strong enough to lead to saturation. Third, we find that a basally weak promoter with high responsiveness, when paired with a strong enhancer, can achieve higher expression than a basally strong but non-responsive promoter (**Fig. 3C-D, 5E**). A simple model in which responsive core promoters can appear nonresponsive due to built-in, proximal enhancers that saturate their capacity for activation^33^ is insufficient to explain our observations. Fourth, we observe similar promoter responsiveness on plasmids and in-genome (**Fig. S8**), suggesting that permissiveness of the chromatin environment does not play a significant role in determining responsiveness. One framework consistent with these observations is that DNA sequence at core promoters specify the recruitment of distinct conformations or compositions of TFIID that differ in their cooperativity with enhancer-recruited cofactors. Particularly for the majority of promoters which lack classical core promoter elements (*e.g.* TATA, Initiator, DPE), key open questions are whether differences in responsiveness indeed reflect differences in TFIID positioning ^75^ or composition ^76,77^, and whether they correspond to sensitivities to other cofactors ^14,16,28^.

Finally, our results offer guidance on how best to use reporter assays to study transcriptional regulation. Reporter assays have been an indispensable tool for bottom-up genetic and biochemical studies of gene regulation, including the original discovery of the enhancer element. However, their usefulness in the genomic era has been debated. Different assays have yielded different measurements of enhancer and promoter activity, have often shown poor correlation with chromatin signatures of the same elements in endogenous context and pointed to different models of enhancer-promoter compatibility^6,33,45,46,55^. This has led to confusion over whether to attribute the discrepancies to suspected confounders^42,43^ (*e.g.,* reporter design^46^, episomes ^40,78^) or novel observations that shift the classical definition of enhancers (*e.g.*, orientation independence)^79,80^. Our results demonstrate that MPRA measurement concordance between different assay designs, with chromatin signatures, and with CRISPR measurements can be markedly improved by avoiding the technical confounders of transcriptional initiation in the enhancer (upstream MPRA) and enhancer sequences in the reporter transcript (STARR-seq).

Thus, we expect that the downstream MPRA and 5’-end mapped, upstream MPRA assays will be more broadly useful for future studies of enhancer-promoter compatibility, regulatory DNA syntax, effects of genetic variants, and gene therapy vector design.

Our study has certain limitations that highlight opportunities for future work. (i) The enhancer and promoter sequences we tested were designed to span a large span of genes previously tested in endogenous CRISPR experiments but are not exhaustive. Additional patterns of compatibility could be observed with larger sets of enhancers or promoters, different lengths of sequences, or engineered minimal core promoters. (ii) We compared enhancer-promoter compatibility in plasmids and integrated into a single genomic locus (*AAVS1*) and obtained very similar measurements. However, because *AAVS1* is endogenously a moderately transcribed locus, it is possible that additional specificity could be observed in other chromatin contexts or each gene’s native chromatin context. (iii) We studied promoter responsiveness in a single cell line (K562). It will be critical to repeat such studies in additional cell types, for example to determine if the responsiveness of a given promoter varies across cell types or contexts.

Overall, our study resolves competing models of enhancer-promoter compatibility and underscores the active and understudied role that core promoters play in enhancer regulation. Future research directions will include further investigation into the sequence and biochemical basis of promoter responsiveness as well as the incorporation of responsiveness into predictive modeling and re-programming of gene expression.

## Methods

### MPRA construct design

We designed the upstream, downstream, and STARR-seq reporter constructs based on our previous design, ExP STARR-seq, which enabled expression quantification of hundreds of thousands of individual enhancer-promoter pairs^33^. Like ExP STARR-seq, all three reporter constructs produce a short, barcoded transcript that is sequenced to both identify the corresponding enhancer-promoter pair and quantify expression. Other core components of the transcript common between all of our assays and ExP STARR-seq include 5’ and 3’ splice sites preceding a short open reading frame (truncated GFP) and an SV40 polyadenylation signal (PAS) at the end of the transcript. The splice site enables increased reporter expression and a splice-junction targeting PCR primer to specifically amplify reporter cDNA rather than any carried over template DNA.

To minimize potential technical confounders and enable comparison with previous results, our STARR-seq reporter construct is identical to the ExP STARR-seq construct. For the upstream design, the relocated enhancer is separated from the promoter by 77bp similar to other upstream, minimal promoter MPRAs^44^. For the downstream design, the enhancer is buffered by 219bp (derived from a previous background sequence^47^) after the PAS to reduce transcriptional interference.

All three designs are cloned into a reporter vector (pCL056) that allows for the MPRA constructs that can be transcribed in cells as either transient episomes or integrated into the genome at the *AAVS1* locus. Used in previous studies for *AAVS1* locus reporter integration^54^, the vector contains homology arms flanking the reporter construct to enable homology-directed repair to integrate the construct into the *AAVS1* locus. The reporter vector also contains a puromycin resistance marker gene (no promoter, including a 3’ splice site) upstream of the reporter promoter position with a separate PAS that both acts to terminate transcription of the *puro* gene when integrated into the *AAVS1* locus and terminate spurious transcription initiation from the origin of replication which could interfere with reporter signal^81^.

### Enhancer and promoter sequence library design

Overall, we designed 2 libraries of enhancer sequences (“21E” and “203E”) and 3 libraries of promoter sequences (“26P”, “78P”, and “500P”) tested in 3 enhancer-promoter combinations (“21E x 26P”, “203E x 78P”, and “21E x 500”). All sequences are 264bp in length unless taken directly from a previous study that used shorter sequences. Promoter sequences from endogenous promoters were taken, on average, from a range of -240 to +20 relative to the endogenous TSS. All genomic sequences and names of genomic sequences use human genome reference hg19 unless otherwise noted. The library compositions of the 3 enhancer-promoter combination libraries are described below.

To explore potential differences between the 6 reporter assays, we first designed the 21E and 26P libraries to include sequences with varied activities from previous studies using upstream-style and STARR-seq-style reporter assays. In the 21E library, sequences were selected to span the dynamic range of 2 previous large-scale reporter assays: K562 genome-wide lentiMPRA^50^ and ExP STARR-seq^33^. For the 3 negative controls, we included 2 synthetic sequences designed and tested to be inactive from a previous study (“noTFBS” and “CTCF”) ^47^ as well as the least active element (“seq5999”) from the genome-wide lentiMPRA screen. The 4 positive control enhancers are all marked by DNase I hypersensitivity, H3K27ac ChIP-seq signal, and P300 ChIP-seq signal and additionally validated by at least one other orthogonal assay for enhancer activity. They include “chr3:128134842-128135106-enhancer_rc” (strongest enhancer sequence in the ExP STARR-seq assay and ABC-predicted, putative enhancer of *RPN1*), “chrX:48659088-48659352” (CRISPRi-validated enhancer of *HDAC6*^1,5^, and two of the top activating sequences (“seq14780” and “seq10228”) from the genome-wide lentiMPRA screen. In the 26P library, most sequences (16) were selected from constitutively or variably expressed endogenous genes ranging in expression from 0 to 900 TPM in K562 cells. 4 sequences from genomic promoters spanning a range of endogenous expression levels (“10409”, “9443”, “9473”, “9060”) from a previous study that also measured promoter activity at the same *AAVS1* locus in K562 cells^53^ were included for comparison. We included 4 negative control promoter sequences with no expected intrinsic promoter activity including 3 inaccessible chromatin regions measured DNase I hypersensitivity in K562 cells (named with prefix “nonDHS”) and a “noTFBS” synthetic sequence also designed and tested to be inactive in a previous study^47^, similar to the negative control enhancer sequence of the same name. We also included commonly used minimal (minCMV, minP, SCP1) and viral promoters (SV40, CMV).

The 78P and 203E libraries were selected from enhancers and promoters from 7 loci with known enhancer-promoter regulatory pairs. In the 203E library, we included sequences of a few categories:

1. Validated enhancers elements from CRISPRi or genetic experiments ^1,5,60^.
2. Putative enhancer elements that were perturbed with CRISPRi and were scored highly by the ABC model but did not significantly regulate any nearby genes (*e.g.,* MYC-NS1^5^).
3. Regulatory elements that when knocked down via CRISPRi resulted in increased expression of a nearby gene.

Because endogenous enhancers are frequently larger than the 264-bp we aimed to synthesize for these oligo pools, we determined whether to add additional tiles of endogenous enhancer elements on a case-by-case basis depending on the size of the element and whether endogenous accessibility at the element was unimodal. Sequences selected for tiling included approximately 100 endogenous enhancer elements close to 50 candidate target genes. All 21E enhancers were also included. In addition to the 3 negative control enhancers in the 21E library, we included 16 additional sequences (2-3 per locus) originating from inaccessible chromatin regions measured by DNase I hypersensitivity in K562 cells (given “element names” with the prefix “control”). In the 78P library, in addition to promoters of candidate target genes from the 7 loci, we also included (i) all 26 promoters from the 26P library for comparison and (ii) +/-80bp shifts in the window of promoter sequence taken surrounding the TSS for 11 endogenous gene promoters, to test if downstream promoter elements (beyond +20) affect promoter properties in our MPRAs (**Fig. S5E-F**).

We designed the final 21E x 500P library to measure promoter responsiveness on a larger, more diverse set of promoters. The 21E library is the same as the one used for the 21E x 26P library. The 500P library primarily selected promoters from endogenous genes with a range of endogenous expression levels which were previously assayed in ExP STARR-seq and classified into random background and P0/P1/P2 classe^33^. For 7 representative, responsive promoters (FTH1, FADS1, HBG2, FTL, JUNB, PIM2, PRDX2), we inserted 2 copies of the GABPA (ETS factor) consensus motif, centered at −10 and −31 relative to the RefSeq annotated TSS to generate a new promoter sequence. For each original sequence, we did the same with YY1 at +1 relative to the TSS. Consensus motifs were taken from the HOCOMOCO v11 CORE motifs. All promoters in the 26P promoter library were also included for comparison with other libraries.

### Library cloning

All MPRAs are constructed with these 3 common fragments: the enhancer sequence library, the promoter sequence library, and a barcoded, open reading frame (ORF) fragment. The downstream assay includes an additional spacer fragment to further separate the enhancer from the PAS. The fragments are amplified with specific primer sets for each design and cloned into either pCL056^47^ or pJT2 (Addgene 255646, derived from pCL056 and described below) with a single Gibson assembly.

We ordered enhancer and promoter libraries as oligo pools from Twist Bioscience with modal sequence length of 300 nt (264 nt plus pairs of 18 nt adapters on each end for amplification). Oligo pools were resuspended in 10 mM Tris (pH 8) and diluted to a working concentration of 2 ng/uL. We then PCR amplified the enhancer and promoter libraries in either a single PCR or 2 sequential PCRs to add Gibson overhangs with respective primers for the upstream, downstream, and STARR-seq design constructs.

For the upstream design, the enhancer library was first amplified in a 50 ul reaction with 1 ul of 2 ng/ul library, 2.5 uL each of 10 uM primers (oJT_ExP_1 and P4_RC), and 25 uL NEBNext Ultra II Q5 Master Mix (Ultra II Q5 MM) (NEB, M0544) with the following thermocycling protocol: 98 C for 30 s, then 4 cycles of 98 C for 10 s, 59 C for 30 s, 72 C for 20 s, then 14 cycles of 98 C for 10 s, 72 C for 20 s, and then a final step at 72 C for 2 mins. PCR products were cleaned with a 1X AmpureXP bead clean and eluted in 20 uL H_2_O. Cleaned PCR product was amplified a second time in a 20 ul reaction with 1 uL of cleaned product, 1 uL each of 10uM primers (oJT_ExP_2 and JR43), and 10 uL of Ultra II Q5 with the following protocol: 95 C for 30 s, then 6 cycles of 98 C for 10 s, 59 C for 30 s, 72 C for 20 s, and then a final step at 72 C for 2 mins.

PCR products were again cleaned with a 1X AmpureXP bead clean and eluted in 20 uL H_2_O. The promoter library was amplified only a single time in a 50 ul reaction with 1 uL of 2 ng/uL library, 2.5 uL each of 10 uM primers (JR44-prom-Fwd and JR45-prom-Rev), and 25 uL Ultra II Q5 with the following protocol: 98 C for 30 s, then 4 cycles of 98 C for 10 s, 62 C for 30 s, 72 C for 20 s, then 14 cycles of 98 C for 10 s, 72 C for 20 s, and a final step at 72 C for 2 mins. PCR products were cleaned with 1X AmpureXP bead clean and eluted in 20 uL H_2_O.

For the STARR-seq design, the enhancer library was similarly amplified in 2 sequential PCRs with identical PCR reaction setups, thermal cycling conditions, and post-reaction clean-up as for the upstream design except for the following changes: primers for the first PCR are DB46 and DB47, primers for the second PCR are oJT_ExPstarr_3 and oJT_ExPstarr_4. Both use an annealing temperature of 59 C. The promoter library was also amplified in 2 sequential PCRs with identical reaction setups, thermal cycling conditions, and post-reaction clean-ups as the enhancer library except for the following changes: primers for the first PCR are oJT_ExPds1_1 and oJT_ExPds_12, and the primers for the second PCR are oJT_ExP_2 and oJT_ExPds_13. The annealing temperature for both PCRs is 62 C.

For the downstream design, both the enhancer and promoter libraries were amplified in a single PCR reaction. The enhancer library was amplified identically to the first PCR of the enhancer library in an upstream design with the exceptions of using primers DB46 and oJT_ExPds_E_rev using a 59 C annealing temperature. The promoter library was amplified similarly in a single PCR with primers oJT_ExPds_P_fw and oJT_ExPds_P_rev with an annealing temperature of 62 C.

The ORF fragment was amplified from the human STARR-seq screen vector (hSTARR-seq_SCP1 vector_block 4, Addgene, #99319) in 2 sequential PCR steps to first introduce a 16 nt barcode with the minimal number of PCR cycles followed by adding Gibson overhangs. For all MPRA designs, the STARR-seq screen vector was first amplified in a 50 uL reaction with 1 uL of 1 ng/uL vector, 2.5 uL each of forward and reverse primers, and 25 uL of Ultra II Q5 with the following general protocol: 98 C for 30 s, then 4 cycles of 98 C for 10 s, 65 C for 30 s, 72 C for 20 s, and a final step at 72 C for 2 mins. Upstream used JR46v2 and oJT_ExP_3, STARR-seq used primers JR46v2 and oJT_ExPstarr_1N, and the downstream design used primers JR46v2 and oJT_ExPds_6N. PCR products were cleaned with a 1X AmpureXP bead clean and eluted in 20 uL H_2_O. PCR products were then amplified in a 20 uL reaction with 1 uL of cleaned product, 1 uL each of 10 uM forward and reverse primers, and 10 uL of Ultra II Q5 with the following conditions: 98 C for 30 s, then 16 cycles of 98 C for 10 s, 30 s at the respective annealing temperature, 72 C for 20 s, and a final step at 72 C for 2 mins. Upstream used JR46v2 and oJT_ExP_3 with a 70 C annealing temperature, STARR-seq used primers JR46v2 and oJT_ExPstarr_2 with a 63 C annealing temperature, and the downstream design used primers JR46v2 and oJT_ExP_4 with a 67 C annealing temperature. PCR products were again cleaned with a 1X AmpureXP bead clean and eluted in 20 uL H_2_O.

Fragments were then assembled vectors via Gibson assembly, transformed into *E. coli*, and bottlenecked to the desired complexity.

In the upstream and STARR-seq designs, the amplified enhancer library, promoter library, and ORF fragments are cloned into pJT2 with Gibson assembly. pJT2 is derived from pCL056 by cutting out the original PAS with HincII and NdeI (NEB) and cloning in the new PAS ordered as a gBlock from IDT, gJT_MPRA_pA, via Gibson assembly. For each 20 uL reaction, we used

### 0.05 pmol of each fragment, 0.05 pmol of NsiI-HF and FseI (NEB) cut and gel-extracted pJT2, and 10 uL of NEBuild HiFi DNA Assembly Master Mix (NEB E2621) and incubated at 50 C for 90 mins

In the downstream design, the amplified enhancer library, promoter library, ORF fragment, and additional spacer fragment gBlock from IDT (ds PAS v2.2) are cloned into pCL056 with Gibson assembly. For each 20 uL reaction, we again used 0.05 pmol of each fragment, 0.05 pmol of NsiI-HF and NdeI (NEB) cut and gel-extracted pCL056, and 10 uL of NEBuild HiFi DNA Assembly Master Mix, and incubated at 50 C for 90 mins.

With each assembled pool of vectors, we next transformed 2 uL into 50 uL of NEB Stable Competent *E. coli* cells (C3040H) by thawing cells for 10 mins, heat shocking at 42 C for 30 s, placing on ice for 5 mins, and pipetting in 950 uL of NEB Stable Outgrowth Medium for 60 mins of outgrowth at 30 C. Serial dilutions of 10 uL (at least 10,000X) were then spread onto ampicillin selection plates to assay transformed library complexity. The remainder of each of the serial dilutions and transformed cells were added to 5 mL of LB with 50 ug/mL ampicillin

(Sigma-Aldrich, A5354) and incubated at 30 C for 24 hrs with shaking at 300 r.p.m. Using the plated serial dilutions, we estimated transformed library complexity, selected the desired dilution for the target complexity, and added 500 uL of the corresponding liquid culture to 50 mL of LB with 50 ug/mL ampicillin to incubate at 30 C for overnight with shaking at 300 r.p.m. Plasmid pools were then isolated from cultures using the QIAprep Spin Midiprep kit (Qiagen, 12143).

### Enhancer-promoter barcode dictionary building

For each cloned plasmid pool, we built a dictionary mapping the 16 bp barcode (BC) within the transcribed sequence to enhancer-promoter pairs.

For the upstream and STARR-seq constructs, we amplified a single fragment containing the enhancer, promoter, and barcode. Upstream constructs were first amplified with oJT_ExP_12_P_lib_fw and JR86 in a 50 uL PCR reaction and 1 ng/uL plasmid pool input (98 C for 30 s, 12 cycles of 98 C for 10 s, 71 C for 30 s, 72 C for 30 s), cleaned with a 1X AmpureXP bead clean, and eluted in 20 uL. STARR-seq constructs were amplified and cleaned similarly to the PCR of upstream constructs but with oJT_ExPds_P_lib_fw and oJT_ExPds_9_EnhBC_Rev at an annealing temperature of 71 C for 14 cycles. Sequencing libraries of both constructs were then pooled with 15% PhiX Sequencing Control v3 (Illumina, FC-110-3001). For the smaller 21E x 26P enhancer-promoter libraries, libraries were sequenced with a MiSeq V3 600-cycle kit (Illumina, MS-102-3003) to sequence the enhancer and promoter as paired end 300-cycle reads (sequencing primers oJT_ExP_1_end_fw and DB44 for upstream and oJT_ExPds_1_PromFw1 and DB47 for STARR-seq) and the barcode as the 18-cycle Index1 read (DB45). Note that the two additional cycles of sequencing read past the end of the 16 bp barcode into a constant region as quality control of the sequencing read and barcode cloning. For the 203E x 78P and 21E x 500P libraries cloned in the upstream design, the enhancer-barcode and promoter-barcode pairs were sequenced separately with NextSeq 500/550 Medium Output v2.5 300-cycle kits (Illumina, 20024905) using sequencing paired-end primer sets oJT_ExP_1_enh_fw and DB47 for the enhancer, JR82 and DB44 for the promoter, and both using DB45 for Index1 BC read. Both used the following read lengths: 150 cycle R1, 18 cycle I1, 8 cycle I2, and 142 cycle R2.

For the downstream construct, we separately amplified a promoter-BC amplicon (primers oJT_ExPds_P_lib_fw and JR86) and an enhancer-BC amplicon (JR89 and oJT_ExPds_9) because the longer distance between the enhancer and promoter made a single amplicon too long to cluster efficiently on Illumina sequencing flow cells. Both sequencing libraries are pooled with 15% PhiX Sequencing Control v3 and paired-end sequenced with either MiSeq V2 300-cycle kits (Illumina, MS-102-2002) or NextSeq 500/550 Medium Output v2.5 300-cycle kits.

Promoter-BC libraries used sequencing primers oJT_ExPds_1_PromFw1 and DB44 while enhancer-BC libraries used primers DB46 and DB47. Both use primer DB45 for the Index1 read to sequence the barcode. Both used the following read lengths: 150 cycle R1, 18 cycle I1, 8 cycle I2, and 142 cycle R2.

All processing code is listed under the Code Availability section. Briefly, enhancer and promoter reads were aligned to their respective libraries using bowtie2 v2.5.4. The resulting BAM files were filtered for uniqueness and alignment quality by only keeping the alignment with the best MAPQ score and a manually determined alignment score threshold. We filtered enhancer-promoter-BC tuples for sequencing artifacts by setting a minimum read count threshold and dropping tuples whose barcode differed by a single base from a >5X more abundant tuple with identical enhancer-promoter identity. We also removed ambiguous pairings (more than one enhancer-promote repair for the same barcode). We confirmed that dictionary quality matched expectation by verifying that barcode nearest-neighbor edit distances were unimodally distributed with a median of 3–4.

### Cell culture

All experiments were performed in K562 erythroleukaemia cells (ATCC, CCL-243). Cells were maintained in a humidified incubator at 37 C and 5% CO_2_ in RPMI-1640 (Gibco, 11-875-119) supplemented with 10% heat-inactivated FBS, 2 mM L-glutamine and 100 units/mL streptomycin and 100 mg/mL penicillin. Cell lines were regularly tested for mycoplasma and authenticated through analysis of RNA-seq data.

### Plasmid transfection

Library transfection was performed as described previously^33^. Briefly, we diluted cells 1:2 in fresh medium daily up to 3 days before nucleofection, maintaining a maximum cell density of 600,000 cells per mL. We then nucleofected 10 million K562 cells with 15 ug of the plasmid pool in 100 uL nucleocuvettes with a Lonza 4D-Nucleofector (program T-016) using the SF Cell Line kit (Lonza, V4XC-2012). Cells were harvested after 24 hrs for subsequent library preparation.

### AAVS1 integration

We adapted a previous protocol for library integration into the *AAVS1* locus integration in K562 cells^47^. Cell lines are generated using TALEN-mediated, homology-directed repair by co-transfection with TALEN L and TALEN R (Addgene no. 35431 and no. 35432). We diluted cells 1:2 in fresh medium daily up to 3 days before nucleofection, maintaining a maximum cell density of 600,000 cells per mL. We nucleofected 10 million K562 cells with 15 ug of the plasmid pool as well as 2.75 ug each of TALEN L and TALEN R in 100 uL nucleocuvettes with a Lonza 4D-Nucleofector (program T-016) using the SF Cell Line kit (Lonza, V4XC-2012). For the larger 203E x 78P and 21E x 500P libraries, we pooled 10 such nucleofections to form 100 million cells nucleofected in total. After 48 hours, cells were treated with puromycin antibiotic for 7 days total at a concentration of 1 ug/mL for the first 5 days and 1.5 ug/mL for the last 2 days to select for the cells in which the reporter construct was stably integrated into the *AAVS1* locus. Cells were removed from the puromycin antibiotic and maintained between 100,000 to 1 million cells per mL for another 7 days to allow for cells to recover and non-integrated plasmid to dilute out.

### MPRA library preparation and sequencing

We adapted our previous protocol for reporter assay library preparation^33^. For the plasmid transfections, we split the 10 million transfected cells, washed the cells twice with PBS, and used half of the cells for Monarch gDNA extraction (NEB, T3010S) and half of the cells for total RNA extraction with RNeasy Plus Spin Columns (Qiagen, 74134). For 21E x 26P *AAVS1*-integrated cell lines, we maintained 10 million cells minimum and eventually expanded cells to 40 million total cells before similarly washing and harvesting 4 x 5 million cells each for gDNA extraction and 2 × 10 million cells each for total RNA extraction. For the larger (203E x 78P and 21E x 500P) integrated libraries, we maintained 100 million cells during cell culture and harvested 10 x 5 million cells each for gDNA extraction and 5 × 10 million cells each for total RNA extraction. We then isolated polyA+ mRNA using the Poly(A)Purist MAG kit (Thermo Fisher Scientific, AM1922) and DNase-treated the polyA+ mRNA using TURBO DNase (Thermo Fisher Scientific, AM2238) in 100 uL reactions incubated first at 37 C for 30 min. We then added 2 uL more of TURBO DNase and incubated an additional 15 mins at 37 C. DNase-treated, polyA+ mRNA was purified with Zymo RNA Clean & Concentrator -25 (Zymo, R1017). We reverse transcribed the purified polyA+ mRNA with the SuperScriptIV First-strand Synthesis kit (Thermo Fisher Scientific, 18091050) using the reporter-specific reverse primer DB31. cDNA was cleaned with a 1X AmpureXP bead clean and eluted in 16 uL of water. We selected for the reporter transcript by amplifying the cleaned cDNA using intron-spanning junction primers (DB32 and DB33) with Ultra II Q5 MM in 50 uL reactions (98 C for 45 s, 12 cycles of 98 C for 15 s, 65 for 30 s, 72 for 30 s). Junction PCR (jPCR) products were cleaned with a 1.5X Ampure XP bead clean and eluted in 16 uL. Both gDNA and jPCR products were amplified using Ultra II Q5 MM in 50 uL reactions and the same final set of primers to add sequencing adapters and sample indices (JR89 and JR90). We performed test qPCRs using diluted input to determine the right cycle numbers. gDNA final PCR used 1 ug input while jPCR final PCR used maximum 10ng input. Both sequencing-ready, final PCRs are cleaned with 1.5X Ampure XP bead cleans and eluted in 15 uL of water. Libraries were quantified using the Qubit 1X dsDNA HS Assay Kit and assessed for length using a D1000 HS ScreenTape (Agilent Technologies, 5067–5582).

Libraries were pooled with 15% PhiX Sequencing Control v3 and single-end sequenced with a NextSeq 500/550 High Output v2.5 75-cycle kits (Illumina, 20024906) with 26 cycles on R1 using custom primer DB45 spiked into the default well and 18 cycles on I1 using fully custom primer JR91.

### 5’-end mapping library preparation and sequencing

In order to sequence the 5’-end of reporter transcripts, we modified subsequent steps in the MPRA library preparation after obtaining purified polyA+ mRNA. We reverse transcribed the purified polyA+ mRNA with the SuperScriptIV First-strand Synthesis kit as before but with both DB31 and template switch oligo (TSO) as RT primers. The TSO sequence is identical to that used in Chromium GEM-X Single Cell 3’ v4 assays from 10X Genomics (10X Genomics, 1000957). cDNA was purified with a 1X AmpureXP bead clean and eluted in 16 uL of water. We then performed 3 rounds of PCR with nested reverse primers to selectively amplify full-length reporter transcripts while adding sequencing adapters and sample indices. For each 50 uL PCR1 reaction, we used 10 uL of purified cDNA, 2.5 uL of each 10 uM primer (TSO_PCR1_F and DB33) and 25 uL of Ultra II Q5 MM with the following protocol: 98 C for 30 s, then 15 cycles of 98 C for 10 s, 65 C for 30 s, 72 C for 60 s, and a final step at 72 C for 2 mins. PCR1 product was cleaned with a 1.5X AmpureXP bead clean and eluted in 15 uL. For PCR2, we used 10 uL of cleaned PCR1 product, 2.5 uL of each 10 uM primer (Truseq_partial_Rd1 and JR90T), and 25 uL of Ultra II Q5 MM in 50 uL reactions (98 C for 30 s, 6 cycles of 98 C for 10 s, 69 C for 30 s, 72 C for 60 s, and a final step at 72 C for 2 mins). PCR2 product was cleaned with 1.5X AmpureXP bead clean and eluted in 15 uL. Final PCR3 to add sequencing handles and sample indices used 10 uL of cleaned PCR2 product, 2.5 uL of each 10 uM primer (oJT_ExP_6 and JR86), and 25 uL of Ultra II Q5 MM in 50 uL reactions (98 C for 30 s, 6 cycles of 98 C for 10 s, 69 C for 30 s, 72 C for 60 s, and a final step at 72 C for 2 mins). PCR3 products were cleaned with a 1.5X AmpureXP bead clean and eluted in 15 uL of water. Libraries were quantified using the Qubit 1X dsDNA HS Assay Kit and assessed for length using a D1000 HS ScreenTape (Agilent Technologies, 5067–5582). For 5’-end only sequencing, libraries were then pooled with 20% PhiX Sequencing Control v3 and sequenced with a MiSeq V3 150-cycle kit (Illumina, MS-102-3001) with 125 cycles on R1 using default primers, 18 cycles on I1 using fully custom primer (DB45), and 8 cycles on I2 using default primers. For full transcript, long-read sequencing, libraries were sent to Plasmidsaurus for Premium PCR sequencing by their Oxford Nanopore Technologies platform.

### 5’-end TSS mapping and quantification

All code for processing and quantification of TSSs is available through the repository listed in the Data Availability section. Briefly, 5’-end mapping sequencing reads were first trimmed to remove the TSO sequence from the beginning then keeping only 42 bases past the TSO. R1 reads were then demultiplexed using I1 reads by enhancer-promoter pair for the 21E x 26P library or by promoter (aggregated over enhancers) for the larger 203E x 78P and 21E x 500P libraries. In the latter two libraries, reads where the corresponding enhancer was seq14780 were first removed. Demultiplexed R1 reads were then aligned to the reporter construct custom reference sequence with the corresponding promoter and enhancer sequences inserted at the library positions. We discarded any alignments with greater than an edit distance of 6 relative to the reference sequence and did not consider any enhancer-promoter pairs with <100 aligned reads passing filter or promoters with <1000 aligned reads aggregated across enhancers passing filter. We further adjusted the exact alignment start position quantification to account for the number of trailing Gs added by the template switching mechanism by shifting the start position by at least 3, then shifting further until either the first non-G base in the read or when the G base matches the reference sequence at the corresponding position. Finally, we quantify TSS frequency as the distribution of trailing G-adjusted, alignment start positions.

### Activity of enhancers, promoters, and enhancer-promoter pairs

Expression of each enhancer-promoter pair was computed from the ratio of barcode (BC) count in the RNA to barcode count in the DNA, averaged across BCs. Specifically, after obtaining BC sequencing reads from polyA+ selected RNA and genomic DNA, we matched BC reads to valid BCs in the BC dictionary using matcha^82^ with maximum 1 mismatch allowed. We then normalized counts to sequencing depth and filter out BCs with fewer than 1 count per million (CPM) in the DNA library. We computed the log-expression per BC by taking the log2 ratio of BC CPM in the RNA library to BC CPM in the DNA library. We added a pseudocount of 0.1 CPM to the RNA BC count for log-expression. For each enhancer-promoter pair, we computed the median BC log-expression score across all BCs mapped to the pair in the BC dictionary.

Unlogged expression was computed identically as above, except no RNA pseudocount was added and no log was taken. To re-center negative control expression at zero and facilitate comparison across assays, we scaled log expression from each assay by an intercept term computed as the average log expression score of negative control promoter sequences paired with negative control enhancer sequences. For subsequent analysis, we excluded enhancer-promoter pairs with fewer than 5 barcodes.

From the enhancer-promoter pair expression, we computed basal promoter activity by averaging log expression across pairs where the promoter is paired with negative control enhancers (annotated in **Table S2-S4**).

Enhancer effect (for enhancers not classified as negative controls) was then computed as the difference between the log expression of the enhancer-promoter pair and the basal promoter activity. Enhancer activity is the enhancer effect averaged across all promoters.

In the upstream assays, we define the promoter activity of an enhancer as the barcode expression of the enhancer-promoter pair when paired with a negative control promoter sequence which is not activatable (e.g. noTFBS, *ESM1*).

### Equal compatibility model

The equal compatibility model assumes that all promoters are equally activated, in terms of fold-change, by all enhancers without the need for enhancer-promoter interaction terms. It is equivalent to the multiplicative model that we previously proposed^33^. Here, we model enhancer-promoter pair expression as the product of basal promoter activity times enhancer activity (Expr_i,j_ = *P_basal,i_* x *E_avg,j_*, where Expr_i,j_ is the expression of promoter *i* paired with enhancer *j*).

Operationally, expression, basal promoter activity, and enhancer activity are all computed as described above in log2 units such that log expression is the sum of log basal promoter activity and average log fold-change in expression from enhancer activation.

### Promoter responsiveness model

To fit the per promoter responsiveness scaling factor, *P_resp_*, according to Equation 2, we first computed the *P_basal_* and *E_avg_*, in log units for each enhancer-promoter library, as above. Then, per promoter, we computed *P_resp_* as the slope of the best-fit line with fixed intercept at y = 0 when regressing the per promoter enhancer effects (log2 units) against enhancer activity (log2 units). The Promoter Responsiveness model-predicted log expression is then computed as the log2 basal promoter activity plus the product of enhancer activity and *P_resp_*. As the fitting of *P_resp_* depends on the enhancer activity computed across promoters in the library, *P_resp_* computed for each library of enhancer-promoter pairs differs based on the dynamic range of enhancer strengths in the enhancer library as well as the promoters with which enhancer activities are computed. Enhancer strength dynamic range was consistent across enhancer libraries as the common set of 21 enhancers we included in all enhancer libraries already contained the two strongest enhancers measured in all assays as well as several negative control enhancers.

Thus, to unify *P_resp_* across the 203E x 78P and 21E x 500P libraries, we accounted for different libraries of promoters by independently computing *P_resp_* with the respective libraries, then scaled *P_resp_* between the two libraries with the slope of the best-fit line for *P_resp_* values of the set of common promoters between the two libraries.

### Comparison of CRISPR-derived enhancer effects with promoter responsiveness

We compared the number and effect sizes of genomic enhancers regulating genes with differentially responsive promoters in our MPRAs by analyzing CRISPRi tiling screens from previous studies that perturbed all DNase-accessible sites around selected genes^1,5^. CRISPR knockdown effect sizes and pre-computed ABC score components were obtained from Supplementary Table 6a of Fulco et al^1^. For genes measured both in our MPRA and comprehensively tiled in CRISPRi screens, we correlated promoter responsiveness with the number of non-promoter, non-CTCF peak, DNase-accessible sites for which perturbation led to a significant reduction in gene expression. For differentially responsive gene promoters, we also compared the number of such elements per gene to the number of predicted non-CTCF peak elements at CRISPRi tiled loci with ABC scores above the suggested threshold of 0.02 as well as the effect on gene expression measured by FlowFISH.

### Responsiveness-ABC model

The Responsiveness-ABC (R-ABC) model incorporates target promoter responsiveness as an exponent on the Activity term of the ABC model numerator. Because the *P_resp_* factor fit from MRPA data is specific to the scale of MPRA-measured enhancer activities, we first confirmed that the MPRA-measured enhancer activity (units of fold change) of the most active 264 bp enhancer sequence tile and the Activity score in ABC (units of RPM) of the correspo16nding 500 bp or larger genomic region were linearly correlated. Then, to exponentiate Activity scores computed in the ABC model, we linearly scaled all Activity scores to the MPRA-measured enhancer activity range before applying the target gene’s promoter responsiveness and afterwards mapped the result back into ABC Activity scale. This Activity score scaled by MPRA-measured promoter responsiveness is then multiplied by the normalized 3D contact value output by the ABC model from input Hi-C data to obtain the final R-ABC score.

### Expression and H3K27ac variability across ENCODE cell lines

To compare regulatory environment variability with expression variability for genes with differential promoter responsiveness, we downloaded uniformly processed RNA-seq and H3K27ac ChIP-seq data for 5 ENCODE cell lines of diverse tissue origins (IMR-90, GM12878, K562, HepG2, HCT116). ENCODE accession IDs are listed in **Table S7**. To focus our analysis on gene expression variability driven by local variation in *cis*-regulation rather than epigenetic gene-silencing mechanisms, we first filtered to genes expressed in every cell line (RNA transcripts per million (TPM) > 5). For each gene expressed in all 5 cell lines, we computed gene expression variation across the cell lines as the coefficient of variation (CV) of TPM expression. We used the Python library deepTools^83^ to obtain uniformly scaled H3K27ac ChIP-seq profiles across cell lines by normalizing BAM file coverage (bamCoverage) by sequencing depth (RPGC method) and dividing coverage by the maximum value in each dataset. We defined a gene’s *cis*-regulatory variation across the 5 cell lines as the mean, pairwise Jensen-Shannon distance between the 5 cell lines’ normalized H3K27ac ChIP-seq profile tracks over a 200 kb window centered on the TSS of the gene. Predicted enhancer-gene links for a gene within each cell type are obtained from ENCODE-rE2G^13^ models previously trained on DNase I Hypersensitivity data from the respective cell type and filtered based on the ABC score component feature.

### Promoter sequence analysis with ProCapNet

We downloaded K562 PRO-cap signal tracks, ProCapNet predicted signal tracks, profile contribution scores, and sequence motif instances of 15 recurrent, high-scoring sequence features generated as previously described (ENCODE accession IDs are listed in **Table S7**) ^64^. Briefly, ProCapNet is a convolutional neural network trained to predict PRO-cap transcription initiation profiles from approximately 2 kb sequence input. The contributions of each base in the input sequence to PRO-cap signal were extracted using well-established model interpretation methods and clustered to uncover DNA motifs learned by ProCapNet. For promoters in our MPRA with genomic origin, we annotated each promoter by intersecting promoter fragment coordinates with ProCapNet motif instance annotations and predicted profile contribution scores. To quantify the relative contribution of each annotated motif within the promoter fragment, we obtained normalized profile contribution scores by dividing the profile contribution scores summed over each base of the annotated motif by the summed profile contribution scores over the entire promoter fragment. Among the 15 recurrent and high-scoring sequence features identified in the original ProCapNet study, we merged the Initiator motifs (CA-Inr and TA-Inr) and TATA motifs (TATA and TATATA) for their high degree of similarity. For analyses that categorize promoters by motif contribution, we binarized the contribution of sequence motifs at promoters by thresholding the normalized profile contribution score of each motif at 0.1.

### Biochemical feature enrichment analysis

We correlated promoter responsiveness with various TF and cofactor ChIP-seq data from the ENCODE Project downloaded and processed as described previously^33^. Briefly, a list of human TF and cofactor ChIP-seq narrowpeak files from the ENCODE Project were downloaded (datasets listed in Supplementary Table 11 of Bergman et al.^33^). MPRA-tested enhancer and promoter sequences with corresponding genomic coordinates were annotated with ChIP-seq feature values as either 0 signal for no overlap or the maximum signalValue of an overlapping peak. Multiple experiment accession numbers corresponding to a single TF or factor (arising from experimental replicates or the same experiment from multiple contributing research labs) were all correlated independently but only the most significant correlation by *p*-value was kept for visualization.

## Data Availability

All sequencing data and processed data generated in this study including building the enhancer-promoter-BC look-up tables, quantifying MPRA expression, and 5’-end mapping of reporter transcripts are available through the National Center for Biotechnology Information (NCBI) Gene Expression Omnibus (GSE334082). Plasmids for MPRA construct cloning are available through Addgene and being held for publication. All external epigenomics datasets and ProCapNet models used are available on the ENCODE and FANTOM portals. Data sources are listed in **Table S7**.

## Code Availability

Code for MPRA experiment preprocessing and analysis are available on GitHub at https://github.com/yj-tan/ExP-MPRA.

## Acknowledgements

We thank C. Ludwig from L. Bintu’s Lab for cloning pCL056. We thank T. Jones, E. Jagoda, A. Baskaran, and D. Bergman for help with analyzing ExP STARR-seq data, J. Schaepe for help in *AAVS1* locus integration, A. Thurm, L. Dunkenberger, T. Wang, and M. Khoroshkin for stimulating scientific discussions, A. Thurm, T. Wang, and P. Guckelberger for manuscript feedback, S. Higashino for keeping the Greenleaf laboratory running, and all the members of the laboratories of J.M.E. and W.J.G. for discussions and feedback. J.M.E. acknowledges support for this study from NIH-NHGRI (R35 HG011324; R01 HG014216), the Applebaum Foundation, and the Novo Nordisk Foundation Center for Genomic Mechanisms of Disease (Novo Nordisk Fonden NNF21SA0072102). W.J.G acknowledges support for this study from NIH (U19AI057266, R01HG013317, R01HL171611, DP1HG013599, R01 NS128028, UM1HG012660). Y.T., M.U.S., and B.R.D. acknowledge the support of the NSF Graduate Research Fellowship (DGE-1656518). B.R.D. also acknowledges the support of the Stanford Interdisciplinary Graduate Fellowship affiliated with Stanford Bio-X. M.U.S. also acknowledges a graduate fellowship award from Knight-Hennessy Scholars at Stanford University. We thank Stanford University and the Stanford Research Computing Center for providing computational resources and support as part of the Sherlock High-Performance Compute Cluster. We also thank the Stanford Genomics facility for providing Illumina MiSeq access and support.

## Author Contributions

Y.T., J.M.E., and W.J.G conceived of the study. Y.T. designed the MPRA assays and 5’-end TSS mapping assay with input from J.R. Y.T. designed sequence libraries and conducted and analyzed MPRA experiments with input from B.R.D., J.M.E., and W.J.G. Y.T. and M.U.S. analyzed ENCODE ChIP-seq data. M.U.S. provided ABC and ENCODE-E2G model fits and features. Y.T., J.M.E., and W.J.G. wrote the manuscript with input from all authors. J.M.E. and W.J.G. jointly supervised the work.

## Supplemental Materials

**Figure S1.**
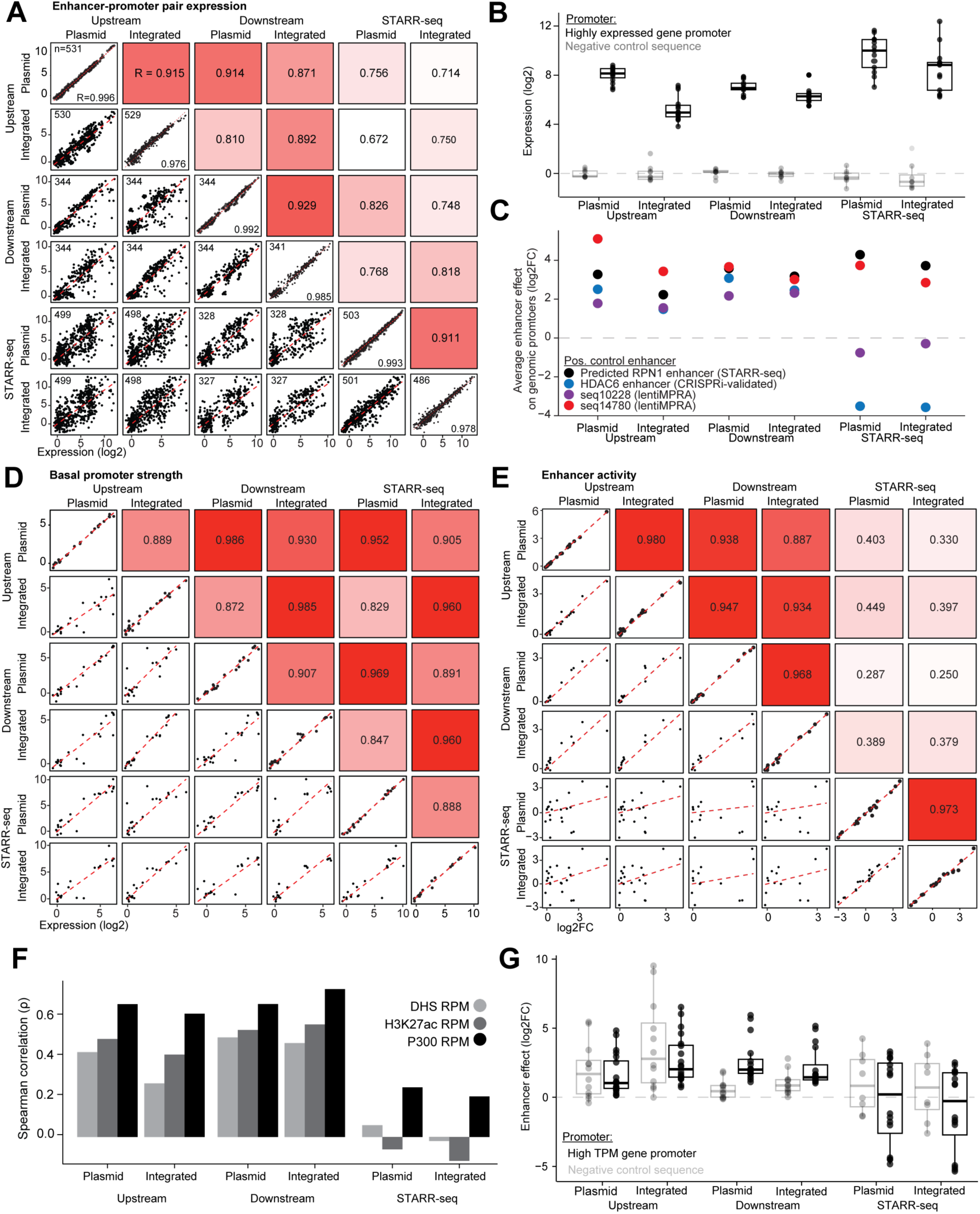
Systematic benchmarking of 6 reporter assays. **A**, Matrix of scatter plots of enhancer-promoter pair expression between all 6 assays tested (bottom triangle, biological triplicate averages). Diagonals scatter two separate biological replicates of the same assay. The upper triangle is shaded according to the Pearson correlation of the corresponding lower triangle scatter plot. **B**, Ability of each assay to distinguish basal promoter activity of 4 negative control promoter sequences from 5 promoter sequences derived from highly expressed genes. Each dot represents one of the promoters paired with one of 3 negative control enhancers. The designation of positive and negative control is described in **Methods**. **C,** Ability of each assay to distinguish the activity of positive control enhancer sequences from negative controls. For each of 4 positive control enhancers in each assay, the log2FC activation averaged over 11 promoters derived from expressed (>10 TPM) genes in K562 is plotted. The assay that previously tested each enhancer sequence is indicated in parentheses. Black dot represents “chr3:128134842-128135106-enhancer_rc” tested in ExP STARR-seq^33^, blue dot represents “chrX:48659088-48659352” (CRISPRi-validated enhancer)^1^, and red and purple represent two of the top activating sequences (“seq14780” and “seq10228”) from a K562 genome-wide lentiMPRA screen^50^ that are also marked by enhancer-associated chromatin signatures. **D**, Same matrix structure as **A**, showing basal promoter activity. Activities are computed per replicate before averaging across replicates. **E**, Same matrix structure as **A**, showing enhancer activity. Activities are computed per replicate before averaging across replicates. **F**, Correlation (spearman) of MPRA-measured enhancer activities (*E_avg_*) with genomic marks of enhancer activity (DHS, H3K27ac ChIP-seq, P300 ChIP-seq) at corresponding native loci. **G**, Boxplots of enhancer effects for 4 positive control enhancers paired with either strong promoters (black) or negative control promoters (gray). Briefly, negative control promoter sequences include designed inactive elements and inaccessible chromatin regions. Positive control enhancers in the 21E library are all marked by DNase I hypersensitivity, H3K27ac ChIP, and P300 ChIP. They are also tested in at least one other direct assay of enhancer activity (reporter assay or CRISPR screen).

**Figure S2.**
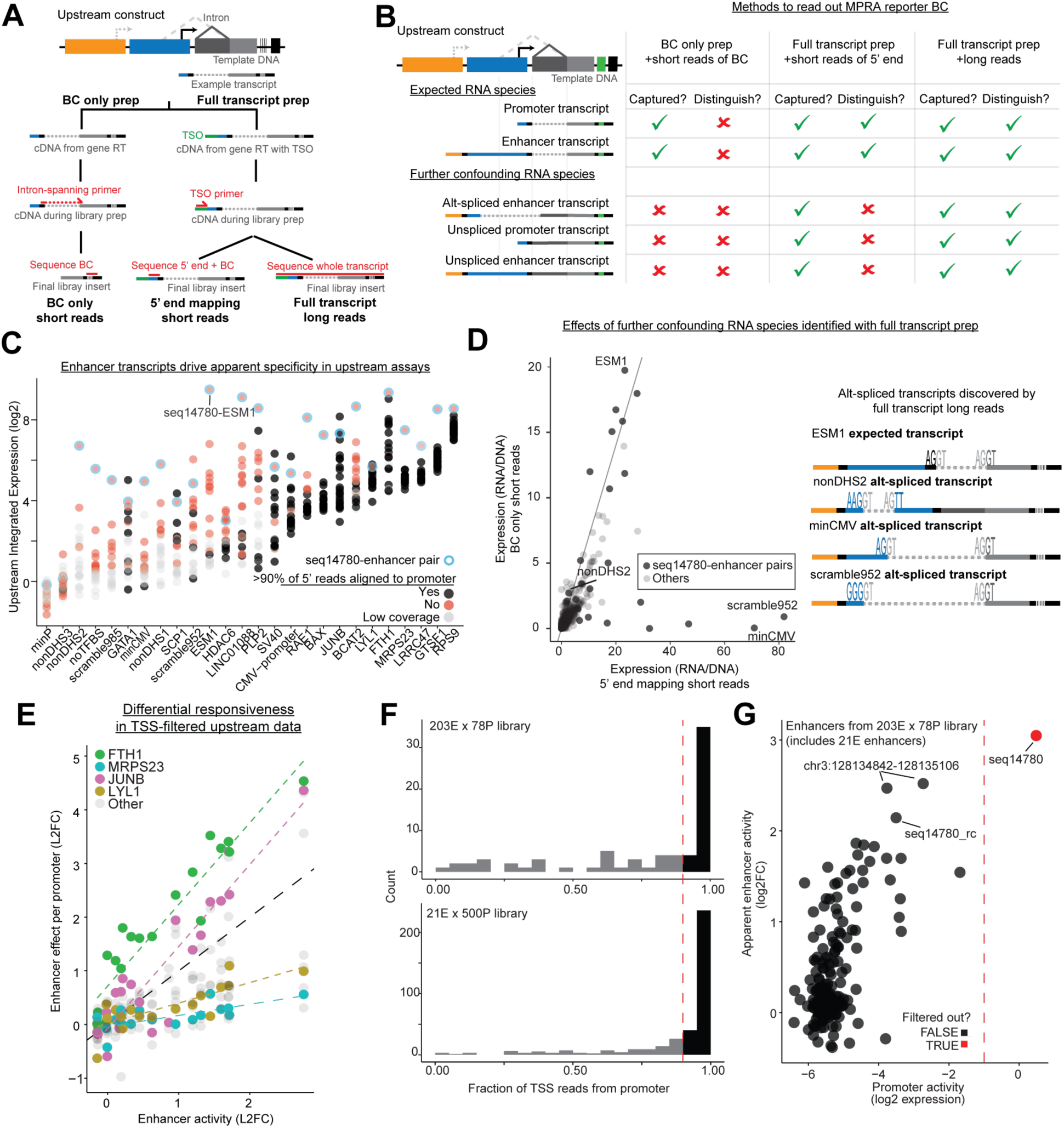
5’-end TSS mapping identifies confounders of and corrects upstream reporter assays. **A**, Schematic of 3 methods reading out reporter expression (2 methods of library preparation and 2 sequencing methods) from an upstream MPRA assay used to identify confounders. “BC only prep” is the typical readout, uses a splice junction-spanning PCR primer to enrich for spliced transcripts, and only reads out the barcode sequence. “Full transcript prep” uses a template switch oligo to add a primer handle to the 5’-end of the cDNA allowing for amplification of the full-length transcript, which can then be sequenced at just the 5’-end or the whole transcript with long-read sequencing. **B**, Schematic of all potential intended and confounding RNA species produced from an upstream reporter design, along with which are both captured in the reporter transcript preparation method (“captured”) and distinguishable from the method of sequencing readout (“distinguish”) by the 3 methods of library preparation and sequencing. For example, the BC only prep “captures” properly spliced transcripts whether they initiate within the promoter or enhancer but cannot distinguish the two types of transcripts because only the BC is sequenced. Alternatively spliced (“alt-spliced”) or unspliced transcripts are rarer but undistinguishable even by full transcript prep with 5’-end sequencing. These species can cause even more unexpected, pathological confounding. **C,** Expression of each E-P pair of the 21E x 26P library in the upstream integrated design where each E-P pair is colored by whether the 5’-end mapping short reads predominantly (>90%) align to the intended promoter TSS (red if yes and black if no). Gray points indicate E-P pairs with too few 5’ TSO PCR reads were detected and/or alignment quality. E-P pairs with the “seq14780” enhancer are outlined in blue. **D**, Expression of E-P pairs in the 21E x 26P library by the BC only short read readout (y-axis) versus the 5’-end mapping short read readout (x-axis). Off-diagonal E-P pairs are due to additional, alternatively spliced reporter transcripts illustrated in **B** that confound expression from the reporter but are undetectable by typical BC only readouts. Examples of a few of the alternatively spliced (“alt-spliced”) transcripts that led to the discrepancy were identified through long read sequencing of the full-length transcripts rather than 5’-end short read sequencing only. **E**, Enhancer effects and enhancer activity computed from upstream integrated assay after filtering out enhancer-promoter pairs confounded by upstream enhancer transcripts. FTH1, MRPS23, JUNB, and LYL1 promoter sequences are genomic in origin, and the endogenous gene is highly expressed. Similar to Fig. 1F and **Fig. S8**. **F**, Histogram the fraction of 5’-end mapping short reads aligning to the promoter for the larger 203E x 78P and 21E x 500P library promoters after first removing seq14780 from the set of enhancer-promoter pairs to analyze. A cutoff of 90% (red dashed line) was used for determining promoter suitable for enhancer activation quantification. **G,** Promoter activity versus apparent enhancer activity of the enhancer sequences in the 203E library. Promoter activity is quantified by the expression of each enhancer with negative control sequence (noTFBS) in the promoter position (BC only short reads). “Apparent” enhancer activity is the average log2FC relative to basal promoter activity of the enhancer over the set of promoters preliminarily kept after filtering in panel **F**, which could still be confounded by autonomous promoter activity of some enhancer sequences. Enhancer sequences “chr3:128134842-128135106-enhancer” and its reverse complement “chr3:128134842-128135106-enhancer_rc” are labeled only with genome coordinates for brevity. Generally, stronger enhancers had more promoter activity. Enhancers with promoter activity greater than -1 were filtered out before further E-P activation quantification.

**Figure S3.**
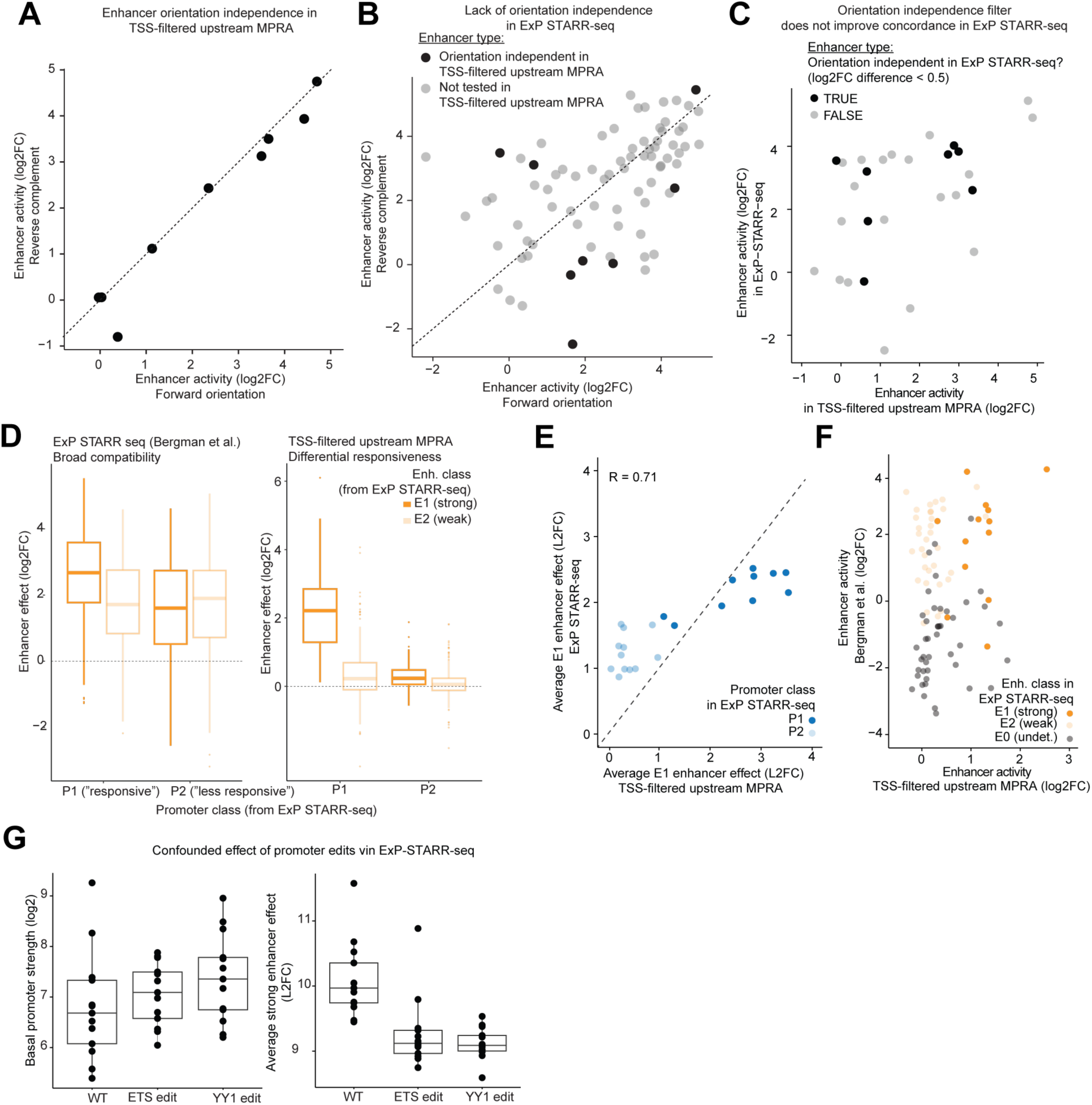
Improved E-P reporter assay corrects previously confounded STARR-seq-measured enhancer activities and E-P compatibility measurements. **A**, Comparison of enhancer activity (averaged over 9 highly responsive promoters) of 8 enhancers tested in both forward (x-axis) and reverse complement (y-axis) orientations (16 total sequences) in the 203E x 78P library in an *AAVS1*-integrated upstream MPRA. **B**, Comparison of enhancer activity in ExP STARR-seq of enhancers tested in both forward (x-axis) and reverse complement (y-axis) orientations averaged over a set of P1 promoters from ExP STARR-seq. Dark black dots are sequences identical to those plotted in **A**. **C,** Comparison of enhancer activities of 28 enhancer sequences tested in both ExP STARR-seq (y-axis) and the 203E x 78P TSS-filtered integrated upstream MPRA (x-axis). The shading of the dots indicates whether the enhancer is orientation independent (solid black) in ExP STARR-seq or not (gray) defined by a difference in enhancer activity between forward and reverse orientations < 0.5 (log2FC). **D**, Comparison of enhancer effects aggregated across previous E1 (strong) or E2 (weak) and P1 (more responsive) or P2 (less responsive) classifications of enhancer and promoter sequences^33^ in both ExP STARR-seq (left) and the TSS-filtered, integrated upstream MPRA (right). **E**, Comparison of the average enhancer effect of E1-classified sequences on P1 (dark blue) vs P2-classified (light blue) promoter sequences tested in both ExP STARR-seq (y-axis) and the TSS-filtered, integrated upstream MPRA (x-axis). Each dot is a promoter tested in both assays. Only enhancer-promoter pairs explicitly tested in both assays were used. **F**, Comparison of enhancer activity in ExP STARR-seq (y-axis) and the TSS-filtered, integrated upstream MPRA (x-axis) for enhancer sequences tested in both assays. Enhancer activities (average log2FC) are computed using promoters in the respective libraries (not required to be common to both assays). Dots represent enhancers and are colored by whether the enhancer was previously classified as E1, E2, or E0. **G,** Box plots mirroring **Figs. 5G** and **5H** with data on WT and ETS/YY1 motif-edited promoters from ExP STARR-seq. Basal promoter activity in ExP STARR-seq was recomputed using sequences annotated as “random BG (background)” in Bergman et al. Strong enhancer log2FC was computed using E1 enhancers in Bergman et al.

**Figure S4.**
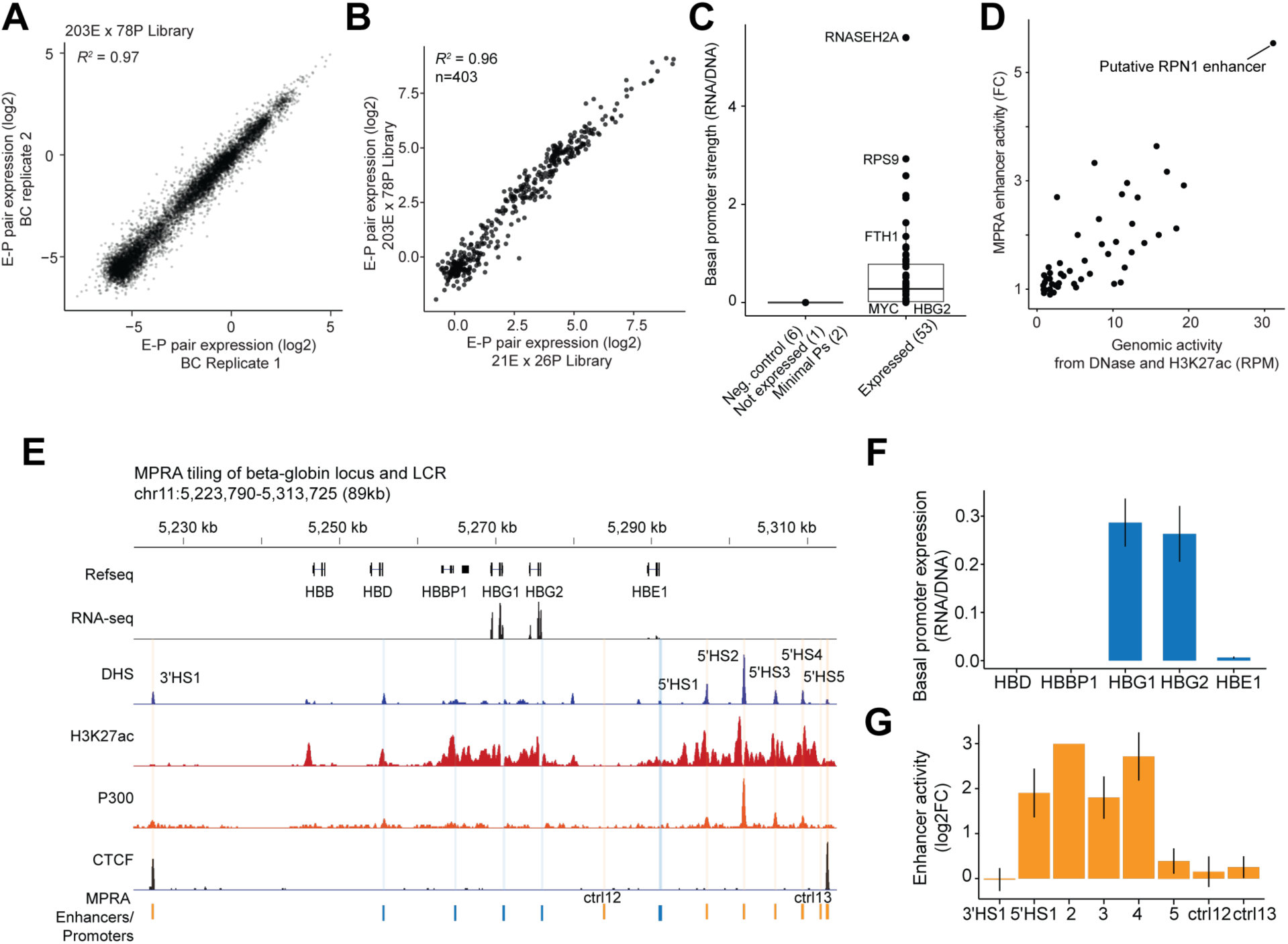
Reproducibility of the 203E x 78P library MPRA and concordance with the corresponding elements at native loci. **A**, Reproducibility of the integrated upstream MPRA with the 203E x 78P library. BC replicate expression is computed per enhancer-promoter pair by splitting the barcodes into two disjoint sets, maintaining at least 5 BCs per set, and separately averaging BC expression (log2). **B**, Scatter plot of enhancer-promoter pair expression for pairs independently tested in both the 203E x 78P library and 21E x 26P library in the integrated upstream MPRA. **C,** Comparison of promoter activity of negative control sequences, ESM1 (which is not expressed in K562), and two minimal promoters (minP and minCMV) which should all have low basal activity versus promoter sequences derived from genes expressed >1 TPM in K562 (“expressed”). **D**, Comparison of enhancer activity from the 203E library after TSS-filtering with the ABC score estimate of activity of the endogenous elements in the genome, computed as the geometric mean of DNase I hypersensitivity and H3K27ac ChIP-seq RPM for putative ABC regulatory elements. For cases of multiple enhancer sequences (264bp) tiling a single >500bp ABC-defined regulatory element, the MPRA enhancer sequence tile with the highest activity was used for this comparison. **E**, The beta-globin locus was one of the 7 loci with well-validated enhancer-promoter regulatory pairs included in this library. Genomic tracks are shown for the 90kb region encompassing the cluster of beta-globin genes and 3’ and 5’ hypersensitivity sites. The cluster of 5’ hypersensitivity sites are collectively called the locus control region (LCR). Shaded columns indicate positions of promoters (blue) and enhancers (orange) within the 203E x 78P library originating from this locus. Sequences “ctrl12” and “ctrl13” tile inaccessible chromatin and are examples of the inaccessible “controls” in the library design. **F**, **G,** Basal promoter activity of the 5 beta-globin gene promoters and enhancer activity of 6 well-validated DNaseI hypersensitivity sitesin the 203E x 78P library, respectively. 5’HS2 through 5’HS5 are abbreviated as “2” to “5”. Concordant with the RNA-seq track in **E**, the HBG1 and HBG2 promoters are expressed at detectable basal levels. 5’HS1-4 have been shown to have enhancer activity and regulate beta globin genes, constituting the enhancers of the beta globin LCR. 3’HS1 and 5’HS5 are accessible and not expected to have intrinsic enhancer activity as accessibility is driven by CTCF-binding. Ctrl12 and ctrl13 (control12 and control13) are also expectedly weak from tiling an inaccessible region.

**Figure S5.**
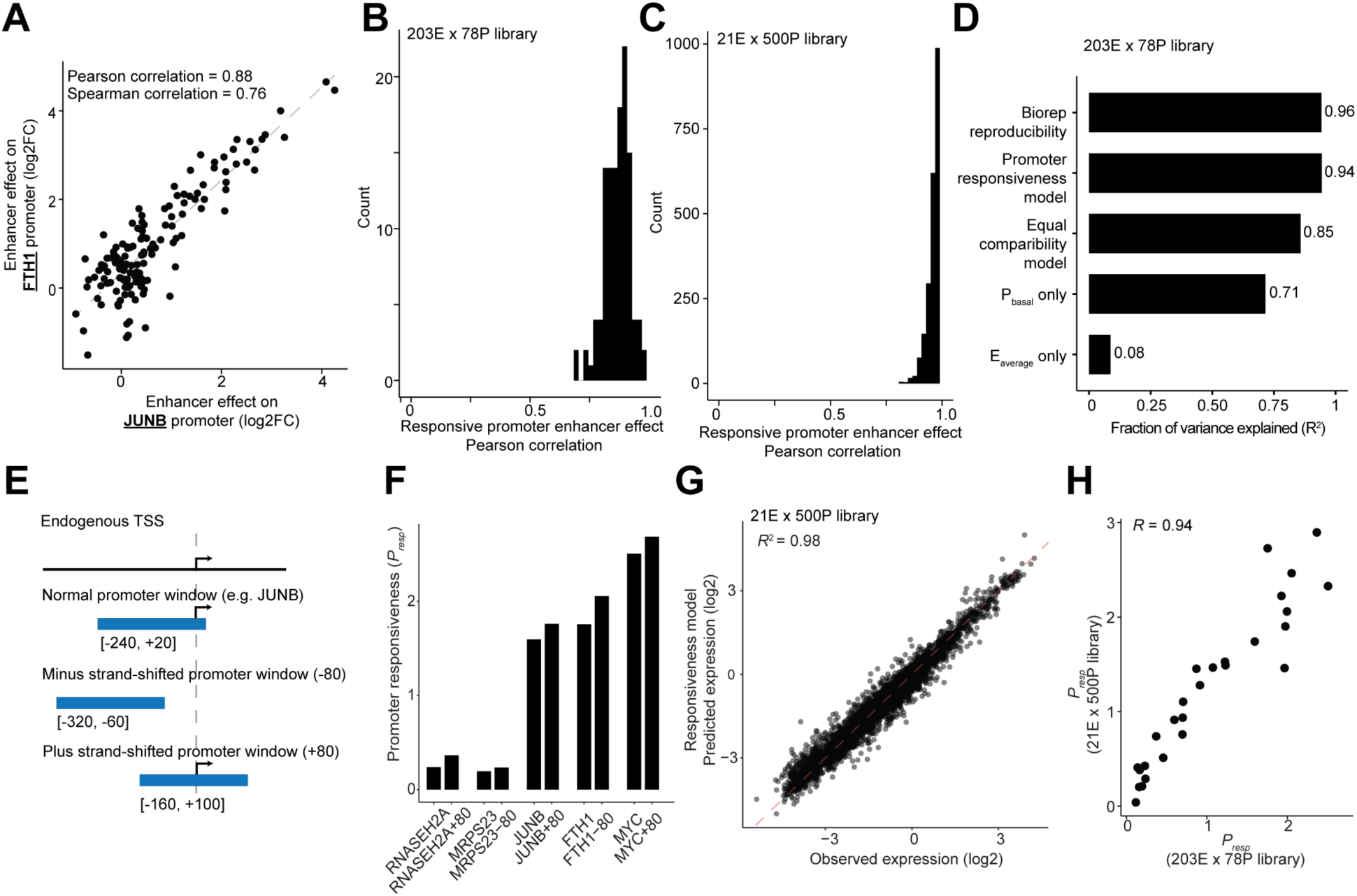
Promoter responsiveness model computed in two separate large-scale MPRA libraries. **A**, Additional example of comparable enhancer effects of many enhancers for two responsive promoters, FTH1 (y-axis) and JUNB (x-axis), in the TSS-filtered, *AAVS1*-integrated, upstream MPRA with the 203E x 78P library. Similar to Fig. 3E. **B**, Histogram of Pearson correlations of enhancer effects between all pairs of responsive promoters identified in the 203E x 78P library. Responsive promoters are defined here as promoters with *P_resp_* > 1. **C,** Same as **B**, except for the 21E x 500P library. **D,** Bar chart comparing the amount of variance explained in 203E x 78P library enhancer-promoter pair expression by different baseline models in comparison with the promoter responsiveness model and biological replicate reproducibility of the experiment (**Fig. S4B**). **E**, Schematic of promoter tiling experiment where different 264bp tiles of various core promoters were tested to see if the precise window around the TSS and sequence further downstream of the TSS affected responsiveness. +/-80bp shifts were defined with respect to the reference genome strand and not the gene’s strand (e.g., an 80bp shift down the positive strand or 80bp shift down the negative strand). For gene promoters located on the positive strand, +80 indicates a shift further towards the gene body while for genes located on the negative strand, this indicates a shift further upstream and away from the gene body. **F**, Comparison of *P_resp_* values of promoter windows shifted about the endogenous TSS. Including additional sequence context downstream of the endogenous TSS does not significantly alter promoter responsiveness. All promoter windows that excluded the endogenous TSS (either +80 or -80 depending on the strand of the gene) were inactive and could not be meaningfully activated by enhancers in the integrated, upstream MPRA. **G**, The responsiveness model explains 98% of expression variance in the 21E x 500P library TSS-filtered, *AAVS1*-integrated upstream assay. Same as Fig. 3G, which was for the 203E x 78P library. **H**, Reproducibility of the fitted responsiveness exponent (*P_resp_*) for promoters tested in both the 21E x 500P (y-axis) and 203E x 78P libraries (x-axis).

**Figure S6.**
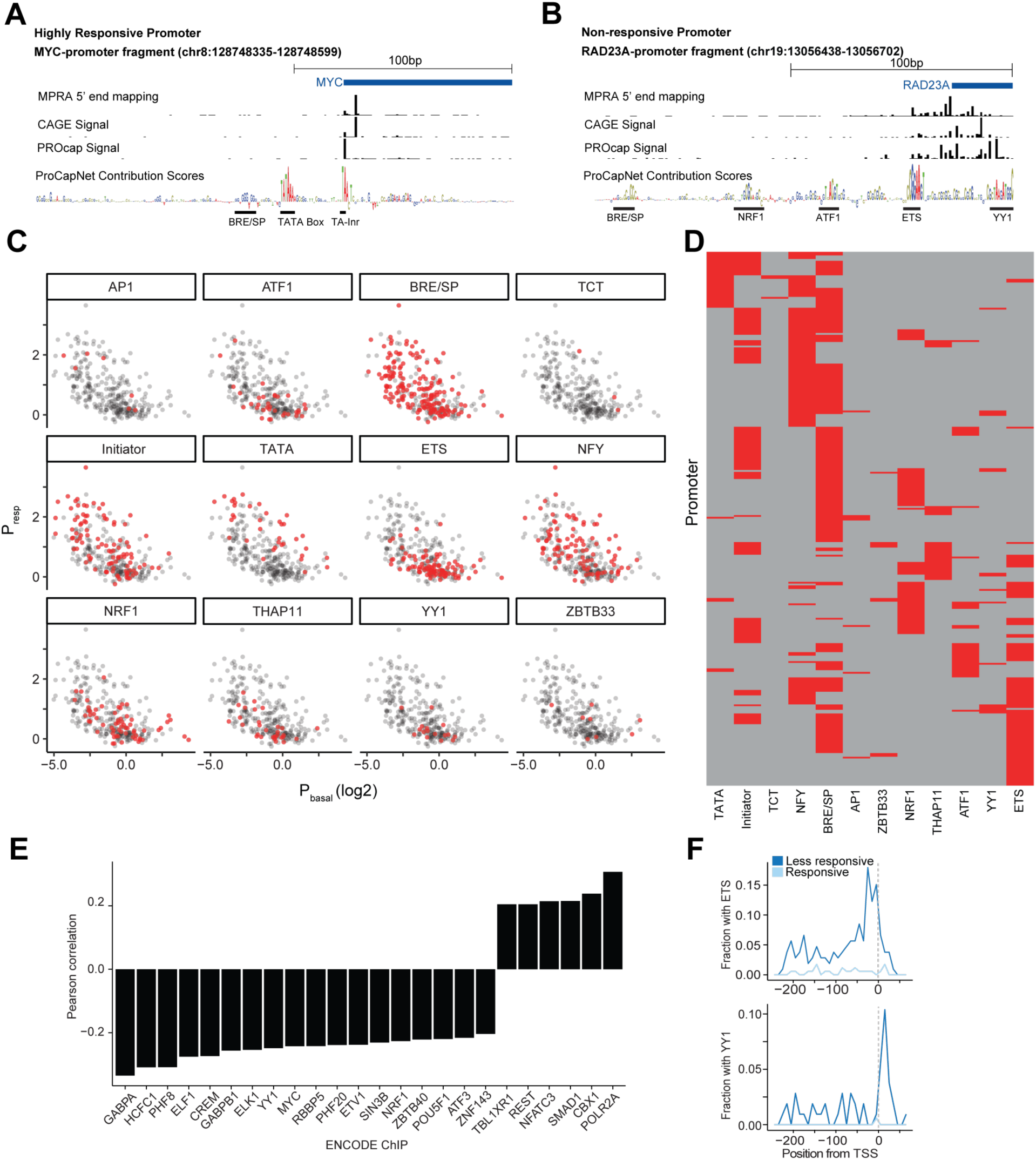
Sequence and biochemical features of promoter responsiveness. **A**, **B**, For a representative responsive gene promoter (MYC) (**A**) and nonresponsive gene promoter (RAD23A) (**B**), respectively, shown genomic tracks compare transcription initiation mapped on our reporters and as mapped endogenously by CAGE and PRO-cap. ProCapNet contribution scores for the corresponding locus are shown at the bottom of each locus, annotating initiation-relevant motifs near each TSS. Genomic coordinates are in hg19. **C**, Promoter responsiveness versus basal promoter activity scatter plot of 500P library promoters (same as Fig. 5A) faceted by the 12 high-scoring, recurrent motifs learned by ProCapNet. Each dot (representing a promoter) is colored red if the respective motif is predicted by ProCapNet to substantially contribute (motif contribution > 0.10) to the initiation signal at the promoter. Motif contribution at the promoter is computed as the sum of the contribution scores intersecting the motif instance divided by the sum of ProCapNet contribution scores over the entire promoter sequence region. **D**, Motifs with substantial contribution (>0.10) in 300 promoters, hierarchically clustered across promoters (y-axis) and sorted by correlation with responsiveness across motifs (x-axis). **E**, Significant Pearson’s correlations of transcription factors and cofactors occupancy at native loci (ENCODE ChIP-seq) with promoter responsiveness. **F**, Frequency of ETS and YY1 motif occurrences normalized by the total number of sequences in responsive (*P_resp_* > 1, light blue) or less responsive (*P_resp_* <= 1, dark blue) promoter classes relative RefSeq annotated TSS’s (gray dotted line).

**Figure S7.**
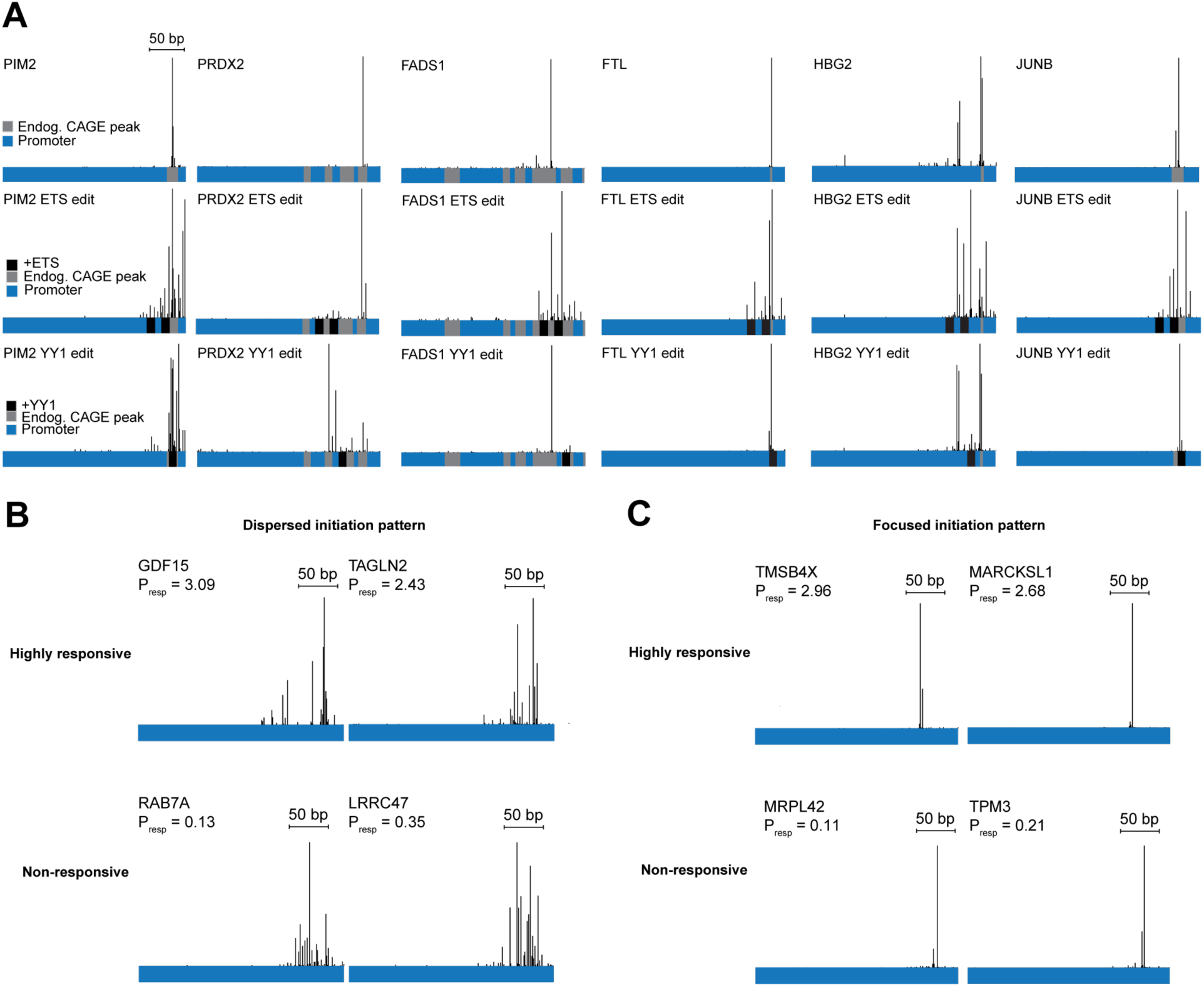
Transcription initiation patterns of sequence-edited and differentially responsive promoters. **A**, Distribution of TSS positions aligned to the promoter region for 6 sets of WT and motif-edited promoters referred to in Fig. 5 measured by 5’-end TSS mapping. The superset of clustered CAGE peak annotations at the corresponding native loci from 1,800 human CAGE samples (gray) and sequence edit locations (black) are indicated. For the ETS and YY1 edited promoters, the CAGE peak annotations for the wild-type sequence are included for reference. For each promoter, TSS mapping reads are aggregated over all enhancers. **B-C**, Both highly responsive (top row) and non-responsive promoters (bottom row) can possess dispersed (**B**) or focused (**C**) initiation patterns. TSSs on the reporter constructs were again measured using the 5’-end mapping assay. Only alignments to the promoter region are shown for simplicity and aggregated over all enhancers.

**Figure S8.**
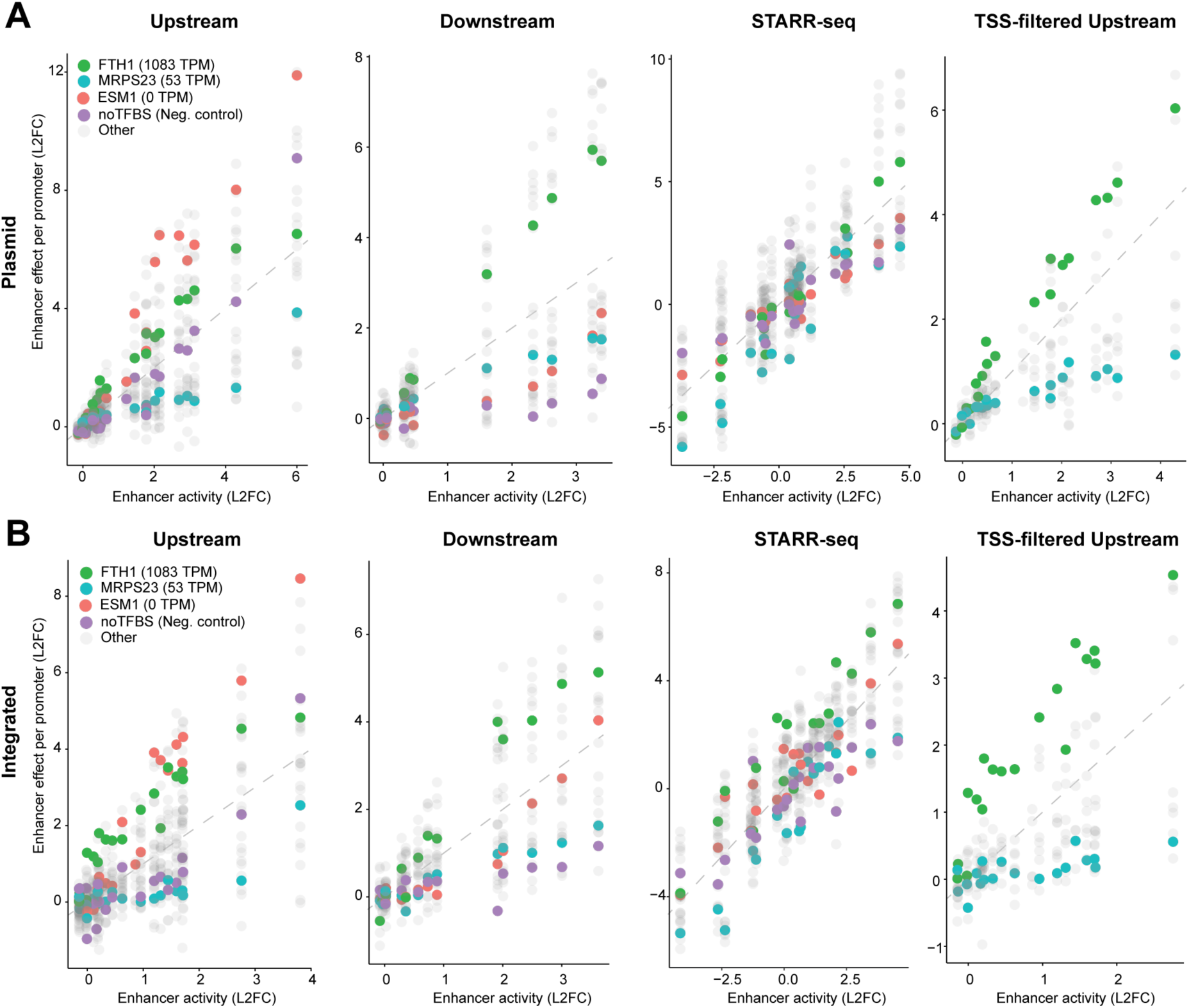
Enhancer promoter compatibility patterns are concordant between plasmid and AAVS1-integrated assays. **A**, Comparison of enhancer-promoter compatibility patterns across MPRA assays (including the TSS-filtered upstream assay) with transiently transfected plasmid templates. Axes are the same as in Fig. 1B and Fig. 1F. Enhancer effects on four promoters representing negative control (noTFBS, purple) and 3 genomic promoters of varying endogenous expression levels in K562 cells (*ESM1* (red), *MRPS23* (blue), *FTH1* (green)) are colored in the upstream, downstream, and STARR-seq assays to highlight qualitative differences in apparent compatibility. noTFBS and *ESM1* promoters are filtered out in the TSS-filtered upstream assay because of confounding enhancer-initiated transcripts. **B**, Same as **A** except with *AAVS1*-integrated versions of the assays.

**Table S1. Primer and oligo sequences.**

**Table S2. Promoter sequences and annotations for the 26P library.**

**Table S3. Promoter sequences and annotations for the 78P library.**

**Table S4. Promoter sequences and annotations for the 500P library.**

**Table S5. Enhancer sequences and annotations for the 21E library.**

**Table S6. Enhancer sequences and annotations for the 203E library.**

**Table S7. External datasets.**

